# Systematic DNA nicking reveals the structural logic of protein recognition

**DOI:** 10.1101/2025.06.30.662289

**Authors:** Yumi Minyi Yao, Michael P. O’Hagan, Karn Onoon, Lihee Givon, Shelly Hamer-Rogotner, Raul Salinas, Naama Kessler, Orly Dym, Tanadet Pipatpolkai, Maria A. Schumacher, Ariel Afek

## Abstract

Transcription factors (TFs) bind to specific genomic sites to regulate gene expression^1,2^. These interactions almost universally require DNA deformation and the accumulation of local mechanical strain within the double helix. As a result, TF-DNA recognition is determined not only by the linear base sequence but also by the spatial alignment of bases and phosphates, as well as their ability to adopt and retain structural deformations^3^.

However, the sequence-centric focus of existing studies makes it challenging to directly probe DNA structural determinants and to decouple their impact from alterations in base sequences, limiting our ability to unravel the key factors influencing binding beyond the sequence identity and leaving significant gaps in our understanding of the principles governing TF-DNA recognition.

Here, we introduce a high-throughput strategy to perturb TF binding sites without altering their base sequence, enabling systematic investigation of the structural features of DNA that govern TF binding. Our method, PIC-NIC, introduces single-strand breaks (SSBs) at every position within the binding site, selectively disrupting backbone continuity while preserving nucleotide identity, with the resulting effects on TF binding measured quantitatively.

Applied to 15 human TFs spanning eight structural classes, and supported by seven high-resolution TF–DNA crystal structures and molecular dynamics simulations, PIC-NIC uncovers discrete backbone positions serving as structural anchor points where nicks can abolish binding, rewire sequence preferences, or even enhance affinity. By decoupling structural and chemical contributions, we demonstrate that DNA mechanics—encoded in backbone geometry and continuity—can independently shape binding specificity beyond the linear code of base identity. These findings shift the paradigm of TF–DNA recognition, establishing the backbone not as a passive scaffold, but as a functional determinant capable of directing regulatory mechanisms through its physical architecture.

## Main

DNA is commonly represented as a linear string of letters corresponding to its base sequence, and transcription factor (TF) binding sites are typically depicted as short sequence motifs. Yet the DNA molecule is far more than a sequence of bases: its ribose sugars, phosphate linkages, and backbone geometry—together with the bases—form an intricate molecular framework that shapes its structural and functional properties^4^. This complexity underlies the precise and nuanced nature of TF recognition: bases provide specific chemical cues, phosphates contribute to electrostatic potential, and the sugar-phosphate backbone supports the spatial arrangement of these elements, sculpting the base-step geometry of the double helix **(Fig. 1a)**^5–7^. Moreover, DNA geometry is inherently dynamic, with certain sequences more capable of adopting and stabilizing strained conformations that are often essential for effective TF binding—establishing DNA mechanics as a fundamental determinant of recognition^8–11^.

**Fig 1.**
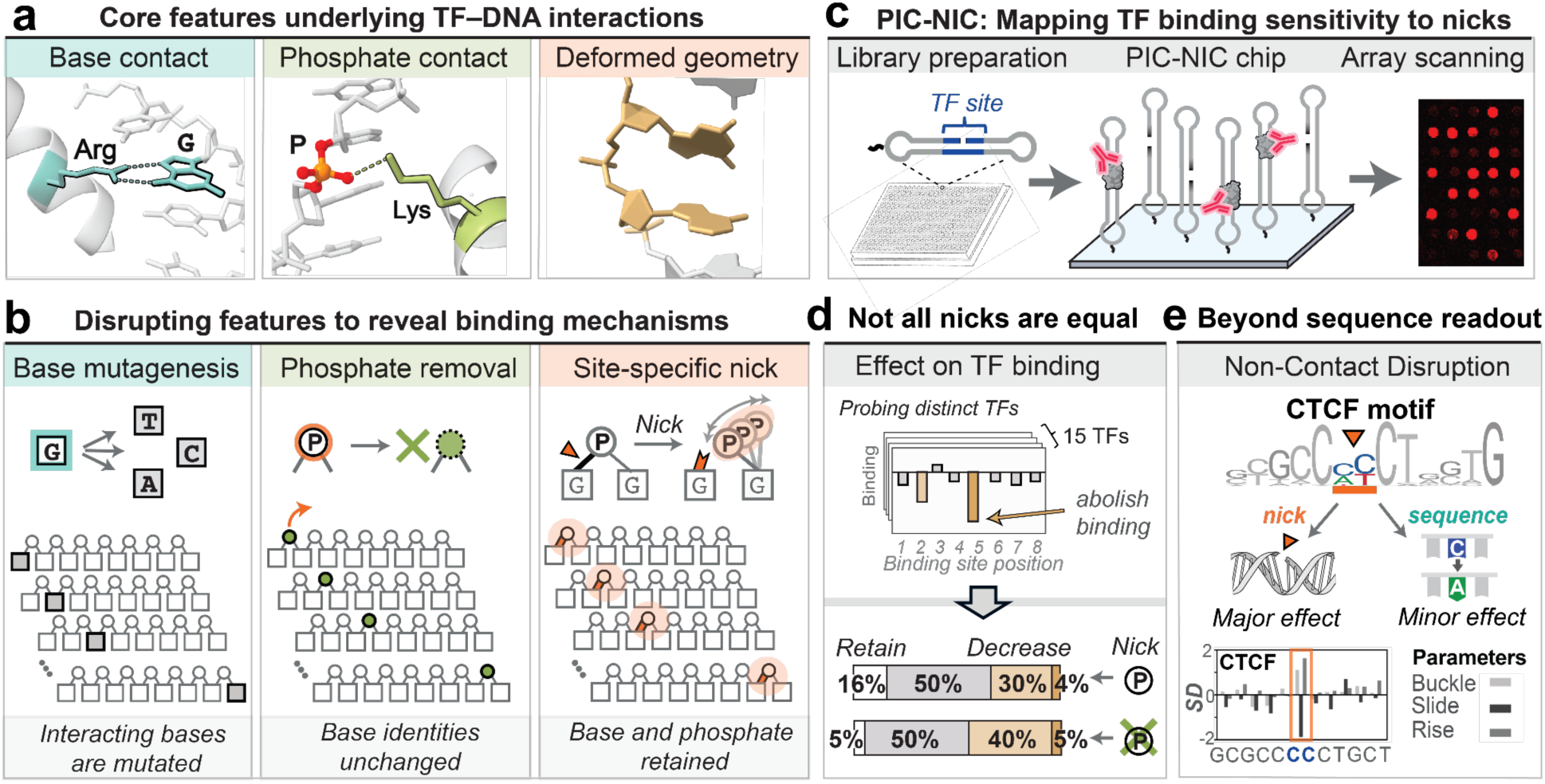
PIC-NIC enables high-throughput dissection of structural contributions to TF–DNA recognition. **(a)** Visualization of key molecular features underlying TF–DNA recognition, using representative complex structures (PDB IDs: 5KKQ and 1QNE). **(b)** Conceptual framework underlying PIC-NIC. Traditional base mutagenesis (left) alters nucleotide identity to probe base-specific contributions. PIC-NIC systematically extends this framework to additional molecular features: phosphate removal (middle), which eliminates backbone phosphates while preserving base identity; and site-specific nicks (right), which preserve both base and phosphate composition while selectively disrupting backbone continuity and geometry. **(c)** Overview of the PIC-NIC platform. Custom-designed nicked DNA libraries are immobilized on microarrays, incubated with target TFs and fluorescently labeled antibodies, and quantitatively scanned to assess binding at each nicked position. **(d)** Schematic depiction of a representative PIC-NIC binding profile (top), illustrating how relative binding levels (y-axis) are measured across all nicked positions (x-axis) compared to the intact control for a given TF. Aggregate analysis across all 15 TFs (bottom; see Methods) classifies nicked positions as retaining strong binding (white), minor disruption (grey), major disruption (gold), or abolishment of binding (brown), for nicks with 5′ phosphate retained (top bar) or removed (bottom bar). **(e)** Disruption at non-contacted positions suggests a role for DNA deformations. As exemplified by CTCF, a nick at a non-contact position (orange) markedly reduces binding, while base mutations at the same position have only minor effects. Structural analysis (PDB ID: 5KKQ) shows that this region exhibits elevated deviations from ideal B-form DNA^9^, with the y-axis representing standard deviations from canonical structural parameters.

Despite this multifaceted nature of TF–DNA recognition, the true role of features beyond direct base readout remains poorly understood^12^. This gap exists primarily because the majority of existing knowledge derives from studies focused on sequence variation, in which structural and electrostatic influences are inextricably linked to base identity, making it challenging to distinguish between the contributions of each recognition mechanism.

A conceptual shift—and new experimental tools—are required to disentangle these mechanisms and reveal the full logic of transcriptional regulation.

### PIC-NIC: Mapping the Impact of DNA Nicks at High Throughput

We reasoned that, just as base mutagenesis has long been used to identify critical nucleotides for TF binding, introducing site-specific disruptions to the DNA backbone—while preserving both base and phosphate identity—could serve as a complementary strategy to pinpoint positions where local structural integrity is essential for recognition (**Fig. 1b**).

Single-strand breaks (SSBs) provided an ideal tool for this purpose. In cells, SSBs are a double-edged sword: they pose a threat to genome stability if left unrepaired^13–15^, yet also play essential physiological roles—for example, in topoisomerase activity, where transient SSBs relieve torsional stress during transcription and replication^16,17^. Inspired by this mechanism, we repurposed SSBs as a high-resolution probe of DNA mechanics, enabling local modulation of backbone geometry without altering base sequence. We hypothesized that if a TF relies on the spatial continuity or rigidity of the DNA backbone at a specific site, introducing a nick that relaxes this constraint would disrupt binding—thereby revealing structurally sensitive positions and decoupling structural contributions from base-specific recognition.

To systematically implement this strategy, we developed PIC-NIC (Protein–DNA Interaction Characterization via Nick-Induced Conformations), a high-throughput method that introduces site-specific SSBs at defined positions within TF binding sites (**Fig. 1c**). By disrupting backbone continuity without altering the base sequence, PIC-NIC enables direct and systematic interrogation of structural effects on TF–DNA recognition, free from confounding changes in nucleotide identity. In addition, we engineered and measured two types of single-strand breaks: one that retains the 5′ phosphate group—preserving chemical moieties required for protein interaction— and one that lacks terminal phosphates while maintaining base composition. This design allows finer resolution in evaluating the specific role of phosphate groups in protein–DNA binding.

In PIC-NIC, libraries of nicked DNA are generated either by self-annealing single-stranded dumbbell oligonucleotides or by hybridizing complementary sequences (Extended Data Fig. 1a), systematically introducing single-strand breaks at every position across known TF binding sites on a DNA microarray (**Fig. 1c**, Extended Data Fig. 1b, Methods). Protein binding signal is measured directly on the chip (Methods)^9,18,19^, yielding highly reproducible results (Extended Data Fig. 1c). Signal intensities on the DNA chip quantitatively reflect binding preferences and have previously been shown to correlate with *K_D_* values measured by orthogonal methods^9,20,21^. For selected targets, we performed in-depth assays to directly measure both thermodynamic and kinetic binding parameters. The high-throughput nature of PIC-NIC enables an unprecedented, systematic dissection of how DNA backbone mechanics influence TF binding across diverse structural families.

### Not All Nicks Are Equal: PIC-NIC Reveals Position-Specific Sensitivity in 15 TFs

We applied PIC-NIC to 15 TFs spanning 8 structural families (Extended Data Fig. 2), systematically quantifying binding to recognition sites in which a single, site-specific nick was introduced at each position across the motif (Extended Data Fig. 3, Supplementary Table 1). While one might anticipate that disrupting backbone continuity at any position would uniformly impair binding, PIC-NIC revealed a strikingly heterogeneous and position-specific response. While many positions tolerated backbone breaks with high levels of retained TF binding, others were highly sensitive to nicking—highlighting discrete sites where DNA backbone integrity is critical for TF recognition.

We parsed the impact of backbone disruption by classifying sites into four categories: retention of binding (<10% decrease in signal), minor disruption (10–50%), major disruption (50–90%), and abolishment of binding (>90%) (Methods). When a nick was introduced with the 5′-phosphate retained, only ∼30% of positions exhibited major disruption, while just ∼4% of positions showed complete abolishment—comparable to non-specific control sites **(Fig. 1d)**. Removal of the 5′-phosphate further increased sensitivity: ∼40% of positions showed major disruption in binding, and ∼5% led to near-complete abolishment. These findings raise key mechanistic questions: what makes certain nicking sites highly disruptive to TF–DNA complex stability only at specific positions, while others are functionally neutral?

We envisaged several mechanisms that may underlie the observed position-specific sensitivity. One likely contributor involves subtle geometric changes introduced by a nick near a TF–DNA contact point—for example, a neighboring phosphate—that may lead to loss of this contact. Although disruption of a single interaction might seem insufficient to explain substantial affinity reductions, we suggest that it can, in some cases, perturb the spatial coordination required for binding. For instance, if a specific interacting phosphate functions as a structural anchor— supporting multiple protein–DNA contacts—its displacement may alter TF docking dynamics and destabilize the entire binding interface.

A second mechanism may involve nicks at sites that are deformed by TF binding. While base pairing and stacking preserve structure in relaxed DNA^22^, strained sites—such as those under bending or torsional stress^23^—may rely more heavily on backbone integrity. A nick at such a site could tip the balance and therefore prevent the DNA from achieving or maintaining its bound conformation.

### Nick-Sensitive Sites Imply Structural Readout

If these mechanisms contribute to the observed sensitivity, we hypothesize that local structural deviations are enriched at nick-sensitive sites. To evaluate the contribution of local DNA structure to TF recognition, we first analyzed a subset of 13 TFs for which high-resolution co-crystal structures with DNA were available in the RCSB Protein Data Bank (PDB). We first focused on sites where the nucleotides flanking the nick do not participate in hydrogen bonding or phosphate interactions with the protein, to minimize the confounding effects of direct contacts.

Remarkably, even in the absence of these direct contacts, 30% of these positions exhibited a significant reduction in binding upon nicking (Supplementary Table 2). Structural analysis of these nick-sensitive, non-contact positions revealed elevated deviations in base-step parameters— particularly enriched in twist and buckle—compared to unaffected sites (Supplementary Table 2). These findings support local DNA conformation as a key determinant of binding affinity at these positions. The enrichment of deviated twist and buckle specifically aligns with the known role of nicks in relieving torsional strain, as observed in the catalytic mechanism of topoisomerases.

Importantly, nick-sensitive sites do not consistently overlap with positions where base mutations strongly impair binding (Extended Data Fig. 3, Supplementary Table 3-4). In certain cases, nicks disrupt binding at positions largely unaffected by base substitutions, while in others nicks have little effect on binding even if the position is highly sensitive to base identity.

For example, when probing the architectural factor CTCF, we found that nicking a specific position where base identity plays a relatively minor role resulted in one of the most pronounced reductions in binding (**Fig. 1e**, top panel). Analysis of the CTCF–DNA complex (PDB ID: 5KKQ) revealed that neither the bases nor the adjacent backbone phosphates at this position form direct hydrogen bonds with the protein, consistent with limited dependence on base identity at this position. Yet the local base-step parameters at this position deviate sharply, with pronounced distortions in buckle, slide, and rise^24^ (**Fig. 1e**, bottom panel).

These observations support a model in which impaired binding arises not from the loss of direct chemical readout, but from disruption of a strain-adapted DNA geometry that facilitates recognition. To further dissect the mechanisms underlying nick sensitivity at sites lacking direct base contacts, we turned to ETS1—an oncogenic TF with rich structural and biochemical characterization^25^.

### Backbone Anchoring Shapes ETS1 Recognition

ETS1 exhibited sharply contrasting, position-dependent responses to single-strand nicks: while some sites tolerated backbone disruption with minimal impact, others showed severe impairment of binding. Strikingly, introducing a nick and removing the 5′ phosphate within the GGA(A/T) core motif—where ETS1 forms direct hydrogen bonds with all three GGA bases^26,27^—had only a minor effect on binding. In contrast, disrupting peripheral positions that engage phosphate contacts (**Fig. 2a**) led to a pronounced loss of binding (**Fig. 2b**, left panels), underscoring the critical role of these peripheral interactions in stabilizing the ETS1–DNA complex.

**Fig 2.**
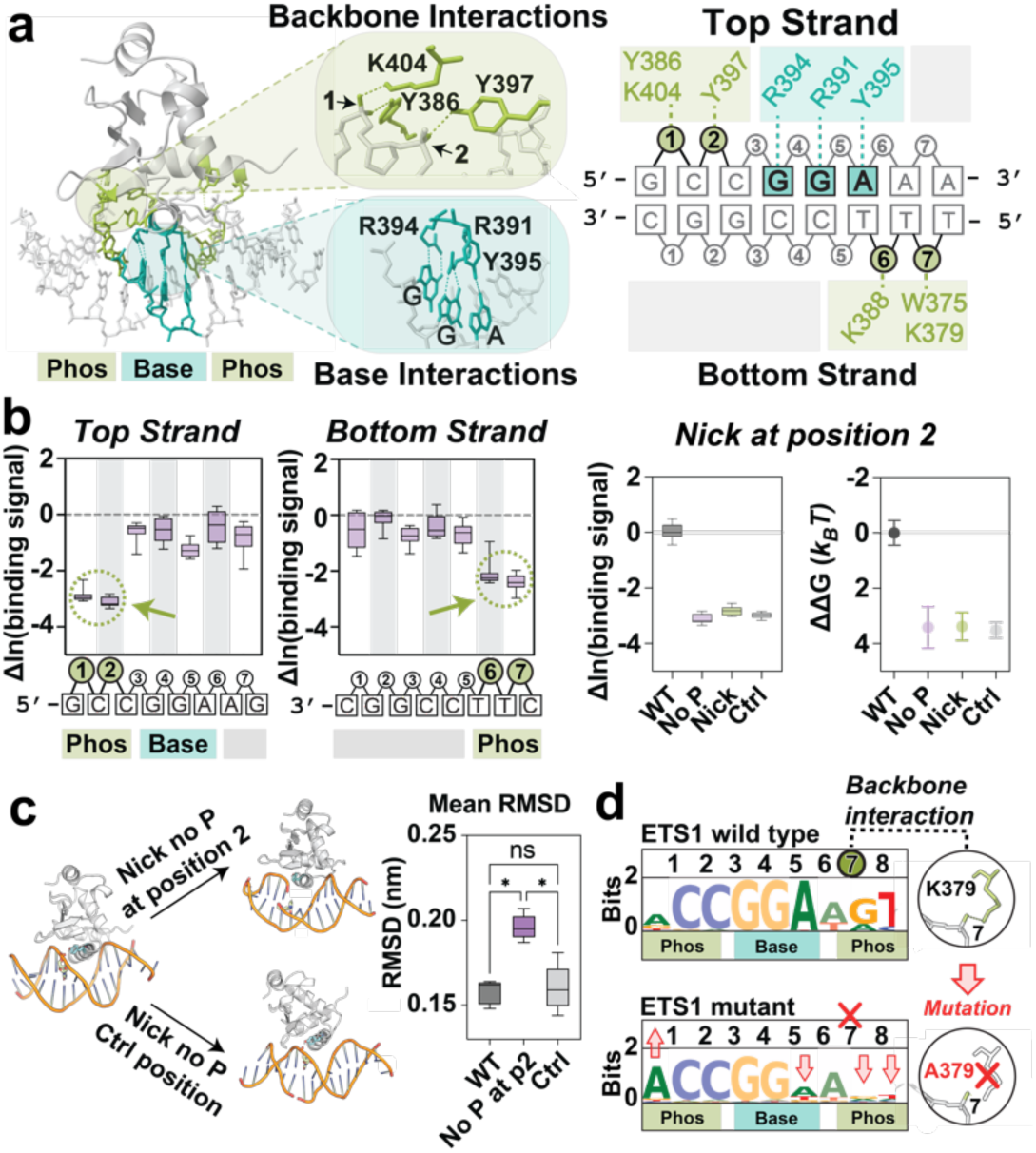
Disruption of phosphate contacts reveals anchoring roles in ETS1–DNA recognition. **(a)** ETS1 forms base-specific and phosphate-specific interactions. In the ETS1–DNA crystal structure (left, PDB ID: 1K79), residues contacting DNA bases are highlighted in cyan, and those contacting backbone phosphates in green. Insets show close-up views of representative phosphate (top) and base (bottom) contacts. Schematic contact maps (right), using consistent color coding, highlight key protein–DNA contacts on both strands, with annotated positions spanning the binding site sequence. **(b)** PIC-NIC phosphate removal profiles (left) show the log difference in binding signal relative to the intact site (y-axis) across positions in the top and bottom strands of the ETS1 binding site (x-axis). The profile reveals that removing phosphates at contact positions markedly reduces ETS1 binding (dashed green circles). Each box in the plots represents 10 replicate measurements. At position 2 (right panels), PIC-NIC and BLI measurements comparing nicks with and without phosphate to the intact sequence reveal that restoring the phosphate does not rescue binding, yielding affinities comparable to a non-binding control. **(c)** Representative snapshots of MD simulations (Methods) reveal reduced complex stability and greater structural disruption at the protein–DNA interface when a nick lacking the 5′ phosphate is introduced at position 2, as evidenced by elevated phosphate backbone RMSD values (5′ GCGGAAA 3′) compared to both the control nicked sequence and the wild-type duplex. Box plots display the average RMSD (compared to the initial frame) over the 1000 ns of simulation for five replicates per DNA/protein complex construct. **(d)** Mutation of a phosphate-contacting residue alters the ETS1 binding profile. PWM logos of wild-type ETS1 (top left) and the K379A mutant (bottom left), which disrupts the phosphate contact at position 7, reveal altered sequence preferences. Red arrows highlight positions with the most pronounced changes in base selectivity, as reflected by shifts in information content (y-axis). Structural insets (right) show the wild-type interaction between K379 and the phosphate at position 7, which is lost in the alanine mutant, eliminating the original hydrogen bond.

Restoring the 5′ phosphate at these disrupted sites partially restored binding at two downstream positions (positions 6 and 7) but failed to restore binding at upstream positions (positions 1 and 2), with position 2 showing the most pronounced loss of binding even when the phosphate was retained (**Fig. 2b**, left panels). Focusing on this site—which also exhibits pronounced twist and buckle deviations (Supplementary Table 2)—we performed Bio-Layer Interferometry (BLI) experiments comparing ETS1 binding affinities and kinetics to the nicked site, the corresponding non-nicked sequence, and a nonspecific control lacking the core motif. These measurements revealed that nicking at position 2 accelerated ETS1 dissociation and reduced binding affinity to the nonspecific level (**Fig. 2b**, right panels, Extended Data Fig. 4a, Supplementary Table 6), providing evidence that this phosphate-contacting site acts as a critical structural anchoring point. To reinforce this observation, we conducted 1 μs x 5 all-atoms Molecular dynamics (MD) simulations: showing that nicking at position 2 induced greater phosphate backbone root mean square deviations at the nicked site and weakened protein–DNA contacts compared to a control nick introduced on the opposite strand, which had minimal impact on binding (**Fig. 2c**, Extended Data Fig. 5).

Recognizing that anchoring relies on coordinated interactions between the DNA and protein, we further probed this mechanism by mutating specific ETS1 residues that directly contact the identified phosphate anchors. To this end, we generated Y397A and K379A ETS1 mutants and performed binding measurements on universal DNA microarrays containing all possible 10-mers^18^ (Supplementary Methods, Supplementary Table 5). Both mutants retained binding specificity, as demonstrated by their position weight matrices (PWMs; **Fig. 2d**, Extended Data Fig. 6a-b) but exhibited altered base preferences compared to wild-type ETS1 (Extended Data Fig. 6). Notably, the K379A mutant, which removes the lysine interacting with the phosphate at position 7, displayed three pronounced shifts in level of base preference across the binding site (**Fig. 2d**). These included reduced selectivity for neighboring bases near the phosphate interaction downstream of the core, as well as a striking reduction in preference for the A within the highly conserved GGA core motif^28,29^—indicating that phosphate contacts directly support base-specific recognition within the core. Additionally, K379A enhanced preference for an A residue at distal sites on the opposite side of the motif, further suggesting that phosphate interactions contribute to long-range architectural coherence across the ETS1–DNA interface.

Together, these findings challenge the traditional view of the DNA phosphate backbone as a merely passive scaffold^5^, suggesting instead that, at least in some cases, it acts as a key structural determinant of TF–DNA binding specificity.

### Nicks Alter TBP Specificity at Hoogsteen-Prone DNA Sites

Beyond simple reductions in binding affinity, we found that TF sensitivity to DNA nicks can manifest in diverse, position-specific ways—including shifts in sequence specificity and even enhanced binding. To explore these possibilities, we focused on TATA-binding protein (TBP) and SOX2, two well-characterized TFs known to induce substantial DNA deformation.

TBP engages the minor groove of the TATA box by inserting two phenylalanine residues at positions 2 and 8, resulting in localized unwinding and a sharp ∼80° bend in the DNA^30^**(Fig. 3a)**. These distortions deviate significantly from canonical B-form DNA and, in certain sequence contexts, are accompanied by the formation of non-canonical Hoogsteen G–C base pairs at the intercalation site (position 8), where guanine adopts the *syn* conformation^31,32^.

**Fig 3.**
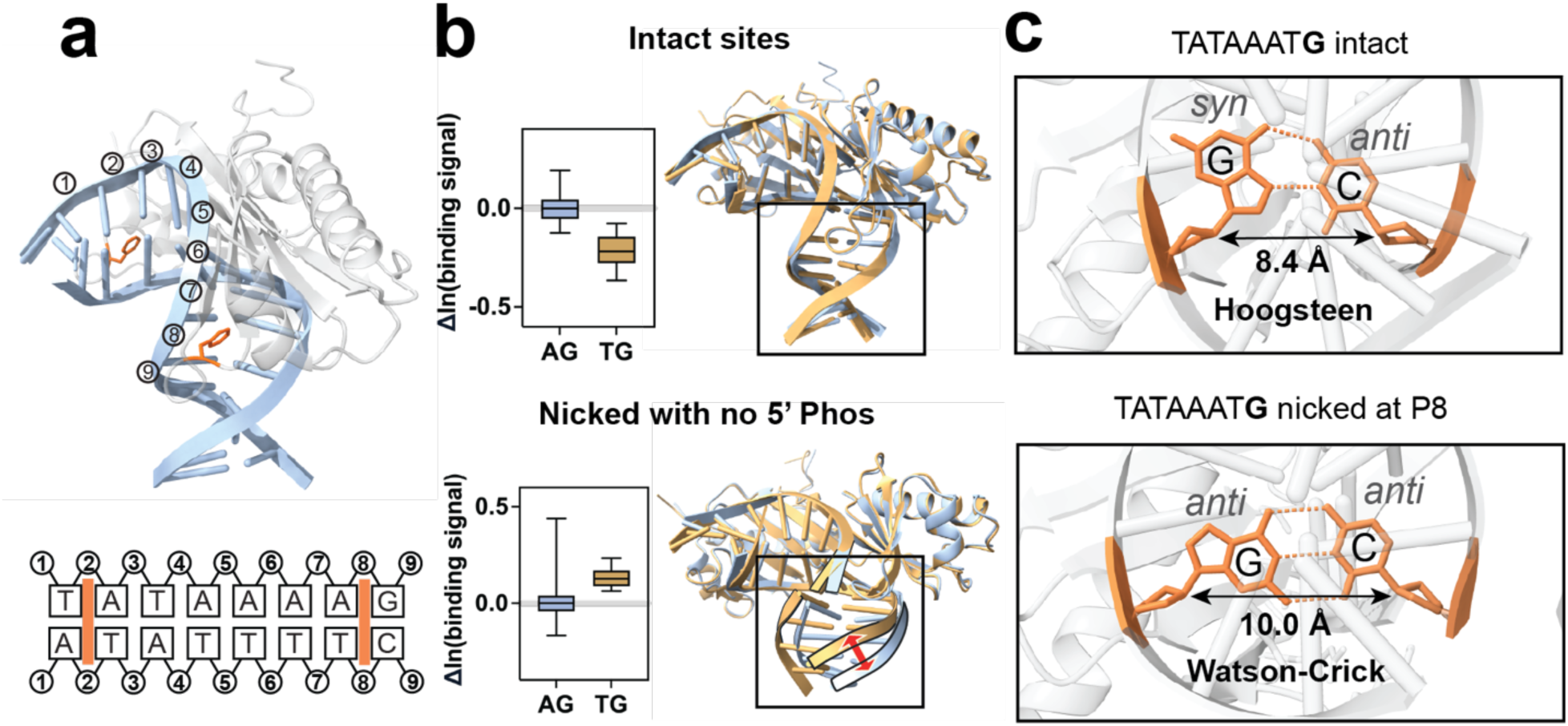
DNA nicks at Hoogsteen-prone positions reshape TBP sequence specificity. **(a)** The crystal structure of TBP bound to a bent TATA-box DNA (PDB ID:1QNE) shows key intercalation positions 2 and 8 (orange) by phenylalanine residues. **(b)** Difference in ln binding signal for TBP to two similar TATA-box variants: TATAAA**AG** (blue) and TATAAA**TG** (gold). In the intact duplex (top), TBP favors the AG variant, whilst upon nicking at position 8 on the bottom strand and removing the 5′ phosphate (bottom), preference switches to the TG variant. Structural overlays of the respective high-resolution crystal structures show high structural similarity in the case of intact DNA (r.m.s.d. = 0.190 Å). In the presence of the nick, increased bending in the nicked TG complex is observed (gold, r.m.s.d. = 0.465 Å; Extended Data Fig.7). **(c)** Close-up base pair geometry at position 8. In the intact TATAAATG complex (top), the terminal G–C of the binding site forms a Hoogsteen pair with an 8.4 Å C1′–C1′ distance. In the nicked complex (bottom), the same base pair adopts canonical Watson–Crick geometry with a widened 10.0 Å distance.

For example, in the TBP–DNA crystal structure (PDB ID: 1QNB) containing the sequence TATAAA**TG** (the **TG** variant), the terminal **G** adopts a Hoogsteen pairing, whereas the closely related sequence TATAAA**AG** (the **AG** variant) retains Watson–Crick geometry in the bound state (PDB ID: 1QNE). Under standard conditions, TBP exhibits higher binding affinity for the Watson– Crick-conforming TATAAA**AG** site **(Fig. 3b)**, possibly reflecting the energetic cost of forming a Hoogsteen base pair^33^.

To directly assess the contribution of DNA geometry at this position, we extracted binding signals for this position from our PIC-NIC dataset and determined four crystal structures of the respective nicked duplexes both with and without a 5′ phosphate (PDB IDs: 9OW8, 9OW7, 9OWZ, and 9OWI; Extended Data Table 1). These experiments revealed several key findings. First, the Hoogsteen pairing observed in the TATAAA**TG** complex reverted to a canonical Watson–Crick geometry upon nicking **(Fig. 3c)**, despite the sequence remaining unchanged. Second, this conformational shift was accompanied by a reversal in binding preference: TBP now exhibited higher affinity for TATAAA**TG** than TATAAA**AG**. These results highlight the key role that DNA conformation can play in influencing TF sequence specificity, where structural changes can convert a previously suboptimal sequence into a preferred binding site under altered geometries

Remarkably, the nicked TATAAA**TG** variant lacking the 5′ phosphate exhibited an even more extreme DNA bend than either of the non-nicked complexes or the nicked TATAAA**AG** variant (**Fig. 3c**, Extended Data Fig.7, Supplementary Table 7). This enhanced deformation suggests that nicks can unlock rare DNA geometries, stabilizing previously inaccessible interactions, and reveal latent sequence preferences by shifting the conformational landscape available for recognition.

### Nick-Facilitated DNA Geometry Accelerates SOX2 Binding

The pioneer TF SOX2 is another well-characterized protein known to induce sharp DNA deformation upon binding in the minor groove **(Fig. 4a)**, inducing a pronounced bend—typically between 70° and 85°—that facilitates chromatin remodeling and transcriptional complex assembly^34,35^. Position 3 of this motif represents an under-twisted site with a minor groove width exceeding 13 Å, more than double the typical 6 Å in B-form DNA, indicating a significant local expansion **(Fig. 4b**, bottom panels). Strikingly, PIC-NIC identified a unique case of nick-promoted binding within our dataset at this position, with a 3-fold increase in binding signal relative to the corresponding intact duplex **(Fig. 4b**, top panel).

**Fig 4.**
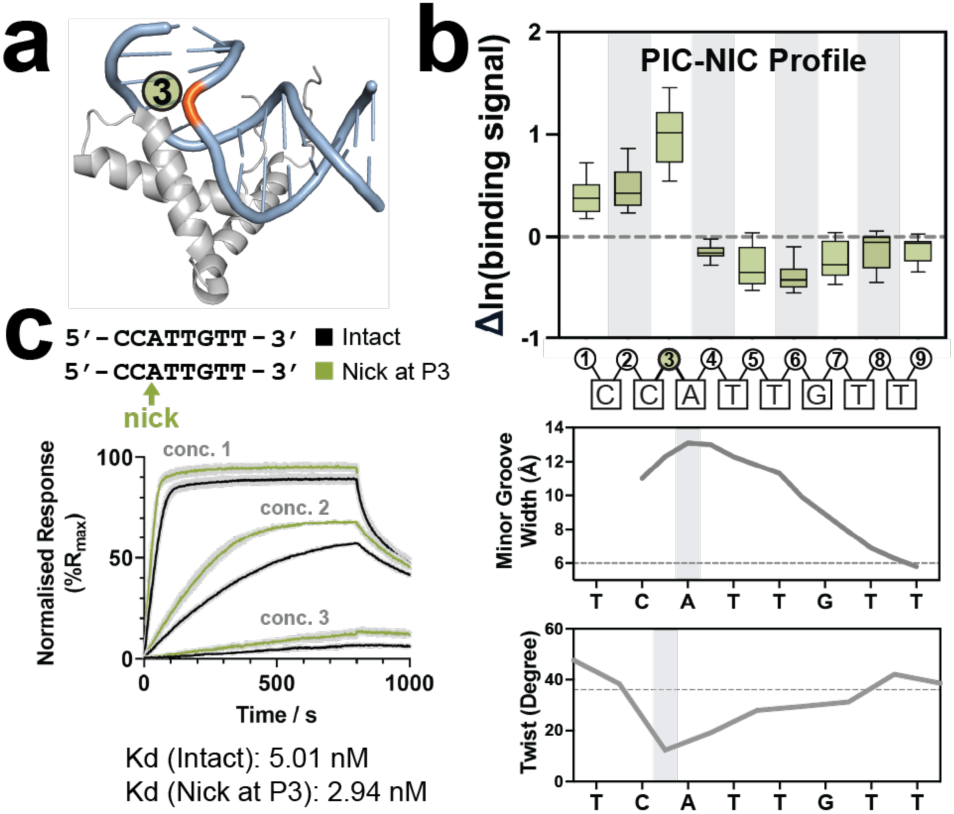
DNA nicks enhance SOX2 binding. **(a)** The crystal structure of SOX2 bound to DNA (PDB ID: 6HT5) shows that position 3 (orange) is highly kinked. **(b)** Top panel: PIC-NIC profile across the SOX2 motif reveals that a nick at position 3 significantly enhances the binding of SOX2. Dashed line represents the binding signal level of SOX2 to intact DNA. Bottom panels: SOX2 structure (PDB ID: 6HT5) reveals an extremly wide minor groove exceeding 13 Å and a significantly decreased twisting angle at this position. **(c)** Bio-Layer Interferometry (BLI) confirms enhanced binding upon nicking at position 3 (green), driven by faster association (*k_on_*).

This observation reveals a distinct mode of sensitivity compared to the previous examples. For example, unlike ETS1, where nicking weakened complex stability and increased the dissociation rate, SOX2 exhibited the opposite trend: BLI measurements indicated that binding enhancement was driven primarily by an increased association rate (**Fig. 4c**, Extended Data Fig. 5b-c). These findings may suggest that the nick partially offsets the mechanical cost required for SOX2 to deform its DNA target, enriching the population of DNA molecules already primed in a recognition-competent conformation, and thereby facilitating more efficient target engagement.

Together with our findings for ETS1 and TBP, these results highlight how TF responses to nicks depend on the local mechanical logic of the DNA and the structural demands of the TF.

These results also raise the broader possibility that other TFs—particularly those acting within compacted or topologically constrained chromatin—may similarly respond to structural perturbations that promote their bound DNA geometries. The structurally sensitive positions identified here may represent sites where deformations—such as those imposed by nucleosomes or partner TFs^36–38^—modulate TF binding without altering the underlying sequence.

### Context-Dependent Roles of Phosphate Contacts in EGR1–DNA Recognition

When all meaningful disruptions are considered—not just the most pronounced—PIC-NIC shows that removing phosphates directly contacted by TFs typically compromises binding, with intact-level affinity retained at only ∼5% of such sites (**Fig. 1d**, bottom bar). This general reduction is expected: the loss of a direct contact should lower affinity by at least the energetic contribution of that interaction, and potentially more if the disruption propagates further. Yet this makes cases where removal of a contacted phosphate has negligible impact—such as at specific positions in the zinc finger protein EGR1—particularly striking.

To investigate this further, we focused on two EGR1 positions where phosphate removal caused the strongest disruption (positions 3 and 6), and compared them to two contrasting cases at position 7, where removal of a directly contacted phosphate on either strand had little to no effect (**Fig. 5a,b**).

**Fig 5.**
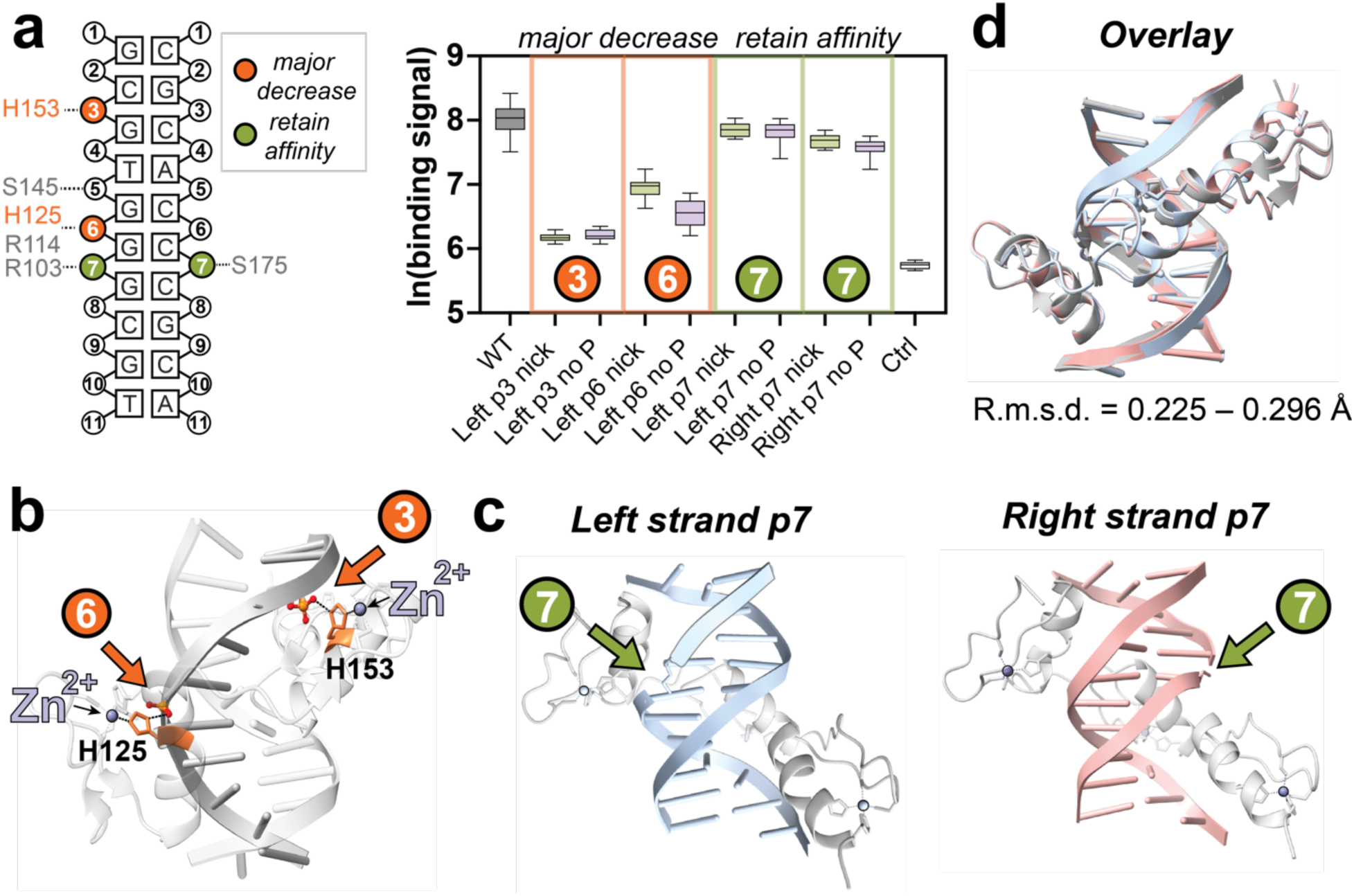
The effects of DNA nicks are context dependent for EGR1. **(a)** EGR1/DNA backbone phosphate contact map for both strands along the binding motif. Orange represents positions where nicks have the most detrimental effect on binding and green represents positions where nicks are tolerated. **(b)** PIC-NIC binding profiles for EGR1 upon introducing site-specific nicks at positions 3 & 6 on the left strand, and 7 on both strands. Loss of the phosphate at positions 3 and 6 strongly reduces EGR1 binding, while both left and right strand nicks at position 7 have little effect, indicating position-specific tolerance to backbone disruption. **(c)** Structural zooms of two zinc coordination sites, highlighting the dual roles of H125 and H153 in DNA phosphate interaction and zinc finger coordination, rendering these positions highly sensitive to nicking. **(d)** High-resolution crystal structures of EGR1 bound to nicked DNA at position 7 on both strands (blue: left strand nick; pink: right strand nick), showing preservation of overall protein–DNA conformation despite the missing phosphate at the break site. **(e)** Superposition of high-resolution crystal structures of EGR1 bound to intact DNA (gray) versus DNA nicked at phosphate position 7 on both strands (blue: left strand nick; pink: right strand nick). Minimal global structural deviation is observed (r.m.s.d. < 0.3 Å, overlay of all atoms), indicating high structural similarity between intact and nicked complexes.

Interestingly, the EGR1 crystal structure (PDB ID: 1AAY, **Fig. 5c**) reveals that the two most disruptive phosphate contacts at positions 3 and 6 involve histidine residues that also coordinate zinc within the C2H2 motif^39,40^. This dual role likely amplifies the functional importance of these interactions, contributing both to local DNA binding and to the structural integrity of the zinc finger domains. In contrast, the phosphate contacts at position 7, which lie between adjacent zinc fingers, appear more dispensable for binding stability.

To further probe this tolerance, we solved two high-resolution crystal structures of EGR1 bound to DNA nicked at position 7 (PDB IDs: 9RIC, 9RJ6; **Fig. 5d**, Extended Data Table 2). In contrast to the TBP–DNA complex, where nicking induced substantial conformational shifts (Extended Data Fig. 7, Supplementary Table 7), the EGR1–nicked structures closely resemble the intact complex, with root mean square deviations (r.m.s.d.) ranging from 0.225 to 0.296 Å (**Fig. 5e**). This may suggest that nicking at these “neutral” positions does not appreciably alter the bound-state conformation. Furthermore, hydrogen bonding analysis shows that beyond the missing phosphate contact, no other interactions are disrupted, and new compensatory interactions may form (Extended Data Fig. 8). These findings support the notion that the energetic contribution of this phosphate, and potentially other non-anchoring phosphates, is modest and can be structurally buffered.

Together, these findings underscore that not all phosphate contacts contribute equally to TF–DNA binding. Some serve as critical architectural elements that integrate local DNA recognition and domain stability, while others are functionally neutral and readily compensated. PIC-NIC enables systematic distinction between these classes, offering nucleotide-level resolution of backbone contributions to recognition energetics.

## Discussion

Our study reveals that DNA backbone integrity—long considered a passive scaffold—plays an active and position-specific role in TF recognition. By systematically introducing site-specific single-strand breaks, we uncoupled DNA backbone integrity from base identity and directly quantified how disruptions in DNA geometry reshape TF binding affinity, specificity, and kinetics across 15 human TFs spanning diverse structural classes. This decoupling exposes a previously hidden layer of recognition logic encoded not in the linear sequence but in the mechanical and geometric architecture of the DNA helix.

We find that discrete backbone positions serve as structural anchor points, where even subtle disruptions destabilize binding, rewire sequence preferences, or in some cases, enhance affinity by reshaping DNA conformation. Notably, these structurally sensitive sites often lie outside the defined base core sequence. For example, peripheral phosphate contacts in ETS1 function as architectural nodes: nicking the DNA or mutating phosphate-interacting residues at these positions triggers long-range destabilization across the interface, altering both affinity and sequence selectivity. These findings challenge the conventional base-centric model of TF recognition and uncover a previously underappreciated layer of structural allostery embedded directly within the DNA backbone.

Beyond single-strand breaks, additional physiological DNA processes—including transcription-induced supercoiling, nucleosome remodeling, chromatin compaction, and binding of other transcription factors—continuously generate local mechanical distortions that reshape DNA conformation without altering base sequence. Our findings suggest that, much like the effects revealed by PIC-NIC, such structural fluctuations may fine-tune TF binding affinity and specificity in vivo, depending on whether they perturb structurally sensitive anchor points. While many deformations are likely neutral, disruptions at these key positions can produce pronounced shifts in TF binding behavior and regulatory output. This adds a dynamic, context-dependent structural layer to transcriptional regulation, where DNA architecture shaped by multiple factors continuously modulates gene expression.

Importantly, our findings establish a structural framework for how DNA damage intersects with both gene regulation and genome stability. Single-strand breaks (SSBs)—the most frequent form of DNA damage—can transiently displace TFs from their binding sites^41^, altering transcriptional programs in a position-dependent manner. While this regulatory consequence is mechanistically clear, its broader physiological relevance in vivo remains to be fully resolved.

Beyond transcriptional modulation, TF binding to damaged DNA can actively interfere with the repair process itself. Growing evidence shows that TFs lingering at damage sites can obstruct repair enzymes, reduce repair efficiency, and elevate mutational risk; conversely, some TFs may stabilize lesion recognition and facilitate repair initiation^42–45^. By systematically charting how backbone discontinuities reshape TF binding at nucleotide resolution, PIC-NIC predicts which lesions will release TFs and open DNA for repair, and which will retain TF occupancy, creating privileged sites where damage may persist and repair is impaired. This framework directly connects the structural logic of TF recognition to repair vulnerability, offering predictive insight into genome stability and mutagenesis.

Together, our findings propose a broader conceptual shift in how regulatory information is encoded in the genome. Beyond the one-dimensional sequence code, we uncover a structural recognition layer embedded in the DNA backbone—a dynamic mechanical grammar shaped by both evolutionary constraints and transient cellular forces, expanding the regulatory logic that governs transcriptional control, genome maintenance, and cellular identity. The PIC-NIC platform can be readily extended to a broader repertoire of transcription factors, offering a scalable framework for systematically dissecting how DNA backbone architecture shapes specificity across diverse regulatory contexts. By illuminating this previously hidden structural layer, PIC-NIC charts a path toward a unified, predictive framework linking DNA mechanics, regulatory specificity, genome stability, and the molecular consequences of DNA damage.

## Methods

### Protein expression and purification

For PIC-NIC experiments, full-length human ETS1, human EGR1 DNA-binding domain (residues 335-423), full-length *Arabidopsis thaliana* TBP, full-length human RUNX1 and human SOX2 DNA-binding domain (residues 19-118) were expressed and purified as described in Supplementary Methods^35,46–49^. Full-length human Max and DNA-binding domain of Myc, Mad and Mnt were chemically synthesized as described previously^50,51^. Human FOXA2 DNA-binding domain (residues 142-269) was expressed by *in vitro* transcription/translation system with PURExpress® *In Vitro* Protein Synthesis Kit (Supplementary Methods)^52^. Full-length human MITF, CREB1, ATF1, SP1 and CTCF were obtained commercially (Supplementary Methods). For ETS1 BLI experiments, murine ETS1 (residues 331-340) was produced as in Supplementary Methods. For SOX2 BLI experiments, the same proteins as in PIC-NIC experiments were used. For EGR1 and TBP X-ray crystallization, the same proteins were used as described above, but with the tag cleaved (Supplementary Methods).

### Universal Protein Binding Microarray

Transcription factor (TF) binding characterization was performed using universal protein binding microarrays (PBMs), as described previously^18,19^. Briefly, commercial microarrays (Agilent Technologies) containing all possible 10-mer or 9-mer sequences were converted to double-stranded DNA via solid-phase primer extension using Thermo Sequenase DNA Polymerase (Cytiva, Catalog #: E79000Y) and a deoxynucleotide triphosphate (dNTP) mixture (dATP, dCTP, dGTP, dTTP). Microarrays were then blocked with 2% (w/v) non-fat dry milk (Sigma, Catalog #: M7409) and incubated with the TF of interest. Binding reactions were carried out in protein-specific buffers (Supplementary Methods).

Pre-incubated protein binding mixtures were applied to individual microarray chambers, incubated for 1 h at room temperature, then subjected to two sequential washing steps. Next, microarrays were incubated for 1 h at room temperature with fluorescently labelled antibody diluted in protein binding buffer supplemented with 2% milk, according to the epitope tag of the protein. Following antibody incubation, microarrays were subjected to two sequential washing steps (Supplementary Methods). Fluorescence signals of the bound proteins were recorded using a GenePix® 4400A scanner. Signal intensities were extracted using GenePix Pro 7.0 software and median pixel intensity was reported for each DNA probe, before further analysis (Supplementary Methods).

### PIC-NIC library design and measurements

PIC-NIC experiments were performed as follows. Libraries of DNA complexes with a nick at every possible position in the TF binding site were designed for each TF, with and without the phosphate at 5′ end of the breakage site. Nick DNA complexes were constructed in two ways. In the first method, a single-stranded oligo was designed to fold on itself upon annealing, forming a dumbbell-shaped DNA, with a nick at the desired position and an amino modifier on the loop of the dumbbell. In the second method, three strands of oligos were designed to anneal and form a duplex, with the two shorter strands being the complements of the longer strand, leaving a nick at the position desired (Extended Data Fig. 1a). The longer strand is labelled with an amino modifier. In the first method, the negative control sequence contained a nick remote from the TF binding site; in the second method, the negative control sequence was the corresponding intact double-stranded duplex.

For each probe, thermodynamic parameters of hybridization were evaluated using the web-based tools from Integrated DNA Technologies and NUPACK, to verify the integrity of annealing under experimental conditions. Simulations incorporated strand concentrations and ionic conditions mimicking experimental settings (150 mM Na⁺, 5 mM Mg²⁺), with appropriate ion correction terms. The maximum complex size was set to 2 for intact duplexes and 3 for nicked complexes involving three strands.

The libraries were spotted onto epoxy-functionalized glass slides using a sciFLEXARRAYER S12 automated non-contact dispensing system (Scienion), followed by rehydration and blocking steps to yield PIC-NIC chips (Supplementary Methods). Protein binding experiments on PIC-NIC chips were performed using the same conditions as universal PBMs (Supplementary Methods). The chips were scanned and the binding signal intensities were recorded the same way as universal PBMs above.

After extracting the measured probe intensities, we quantified the extent of binding disruption at each site by first taking the natural logarithm (ln) of the fluorescence signal. For each nicked site, we computed the percentage signal change relative to an intact control binding site (Supplementary Methods). This was done by calculating the normalized signal change: Signal change = ln(*I*_intact_) – ln(*I*_non-binding_) / ln(*I*_intact_) – ln(*I*_nicked_), where *I*_intact_ is the signal from an intact binding site, *I*_non binding_ is the signal from a non-binding site, and *I*_nicked_ is the signal from the desired nicked site. Sites were then classified into four categories: retention of binding (<10% decrease in signal), minor disruption (10–50%), major disruption (50–90%), and abolishment of binding (>90%).

### Site-Directed Mutagenesis

Site-directed mutagenesis was performed using KAPA HiFi HotStart DNA Polymerase (Roche) according to the manufacturer’s standard protocol. Mutagenic primers were designed (Supplementary Methods) to introduce the desired substitution. Following PCR amplification, the reaction was treated with DpnI (NEB) to digest the template plasmid, and the product was transformed into *E. coli* DH5α. Positive clones were confirmed by miniprep and Sanger sequencing.

### Crystallization and determination of the structure of TBP-nicked DNA and EGR1-nicked DNA complexes

TBP–DNA complexes were prepared and subjected to hanging drop vapor diffusion crystallization screens (Supplementary Methods), resulting in large, well-diffracting crystals suitable for data collection after optimization of initial hits. Data for all crystals were collected at the Advanced Light Source (ALS) on beamlines 5.0.1 and 5.0.2. The data were processed with XDS^53^. The structures were solved by molecular replacement (MR) using a previous structure of TBP (Supplementary Methods) as a search model. MolProbity was used to guide the process of refitting and refinement^54^. See Extended Data Table 1 for the final data collection and refinement statistics for each structure.

EGR-DNA complexes were prepared as previously described^39^ and subjected to hanging drop vapor diffusion crystallization screens (Supplementary Methods), yielding large and well-diffracting crystals suitable for data collection in the initial screens. Data for all the crystals were collected at The Dana and Yossie Hollander Center for Structural Proteomics at the Weizmann Institute of Science. Initial models were iteratively rebuilt and refined using Coot^55^ and Phenix^56^. Model geometry was evaluated using MolProbity^54^. See Extended Data Table 2 for the final data collection and refinement statistics for each structure.

### All-atoms molecular dynamics simulation

The structure of the DNA (5′ TAGTGCCGGAAATGT 3′) with and without ETS1 (PDB ID: 1K79), was nicked at position 7C on either strand using PyMOL, yielding a total of 6 systems. The system was solvated with TIP3P water model and neutralised with 0.15 M KCl using CHARMM-GUI Solution Builder^57^. This resulted in the simulation box dimension of 8.1 × 8.1 × 8.1 nm³. The composition of each simulation box is noted in Supplementary Table 8. The protein was simulated with AMBER99sb-ildn^58^ forcefield and the DNA was simulated with OL21 forcefield^59^. The systems were then energy minimized using a steepest-descent algorithm for 5000 steps and equilibrated for 10 ns with the restraint of 1000 kJ nm^-2^ mol^-1^ on C_α_ of the protein and the phosphate backbone of the DNA. The systems were then equilibrated for 10 ns, maintaining the temperature at 310 K using a V-rescale thermostat^60^ and the c-rescale barostat^61^ for isotropic pressure coupling at 1 bar. Then, the system was simulated using 2 fs timesteps and maintained the temperature at 310 K using a V-rescale thermostat and the c-rescale barostat for semi-isotropic pressure coupling at 1 bar for 1 μs for 3 repeats. All simulations and hydrogen bond analyses were performed using GROMACS 2024.3^62^.

## Data availability

The data that support the findings in this study are available as Supplementary Tables in Excel format. Coordinates and structure factor amplitudes for the TBP-AG-withP, TBP-AG-noP, TBP-TG-withP, TBP-TG-noP, EGR-n7, EGR-ren5P and EGR-ren7 structures have been deposited in the PDB under the accession codes 9OWZ, 9OWI, 9OW8, 9OW7, 9RIC, 9RI6, and 9RJ6, respectively. The PDB entries used in this study are available in Extended Data Figs. 2 and Supplementary Tables 2. Molecular dynamics simulations tpr, mdp and itp files are stored at doi:10.5281/zenodo.15646043.

## Acknowledgements

We thank D. Fass, H.M.A.-Hashim, N. Barkai, H. Hofmann, Y. Levy, V. Salomon, A. Beniaminov, B. & L. Krupkin, and the entire Afek Lab for discussions and comments; S. Albeck, Y. Peleg, I. Miodownik, G. Rosenblum for providing recombinant purified proteins; M. Jbara, R.V. Nithun for generously donating chemically synthesized proteins; E.N. Nikolova for kindly providing expression plasmids; M. Goldsmith for supporting BLI experiments. This work was supported by Israel Science Foundation Grant 1174/22, Prof. Dov and Ziva Rabinovich endowed fund of Structural Biology, Wagner-Braunsberg Family Melanoma Research Fund, the Joel and Mady Dukler Fund for Cancer Research, the Jean-Jacques Brunschwig Fund for the Molecular Genetics of Cancer, the Howard and Janet Rothenberg Pack Endowment Fund for Pancreatic Cancer and Parkinson’s Research, and the Henri Meyer Cancer Endowment (to A.A.), a Nanaline H Duke Endowed Chair and National Institutes of Health grants (R35GM130290 to M.A.S.), and and a Teva-NFBI fellowship award (to Y.M.Y.). We acknowledge Berkeley National Laboratory beamline 5.0.2 and beamline 5.0.1 for X-ray diffraction data collection. The Berkeley Center for Structural Biology is supported by the Howard Hughes Medical Institute, Participating Research Team members, and the National Institutes of Health, National Institute of General Medical Sciences, ALS-ENABLE grant P30 GM124169. The Advanced Light Source is a Department of Energy Office of Science User Facility under Contract No. DE-AC02-05CH11231. We acknowledge beamlines 5.0.1 and 5.0.2 for X-ray diffraction data collection. The Pilatus detector on beamline 5.0.1 was funded under NIH grant S10OD026941. High-performance computing was supported by LKY postdoctoral fellowship (to T. P.), and we acknowledge NTU HPC service for computing support.

## Contributions

Y.M.Y. and A.A. designed and supervised the study. Y.M.Y. developed the assay, generated high-throughput protein-DNA binding data, and analyzed the data. Y.M.Y. performed analysis of crystal structures. Y.M.Y. and M.P.O. contributed experimental data on protein-DNA binding affinities. Y.M.Y. and M.P.O. purified EGR1 recombinant protein. L.G. contributed the plasmids for ETS1 mutants and FOXA2. K.O. and T.P. contributed simulation data. S.H.-R. and O.D. contributed X-ray crystallographic data of EGR1; M.A.S. contributed X-ray crystallographic data of TBP. R.S. purified recombinant TBP for crystallization and N.K. purified ETS1 recombinant protein. Y.M.Y, M.P.O. and A.A. wrote the manuscript, with input from all authors. All the authors critically reviewed the manuscript and approved the final version.

