## Extended Data Figures and Tables for "Systematic DNA nicking reveals the structural logic of protein recognition"

Singapore;

^5^Department of Biochemistry, Duke University School of Medicine, Durham, NC 27710, USA

**This PDF file includes:**

Extended Data Figures 1-8

Extended Data Tables 1-2

**Extended Data Fig. 1 PIC-NIC Pipeline and experimental reproducibility**

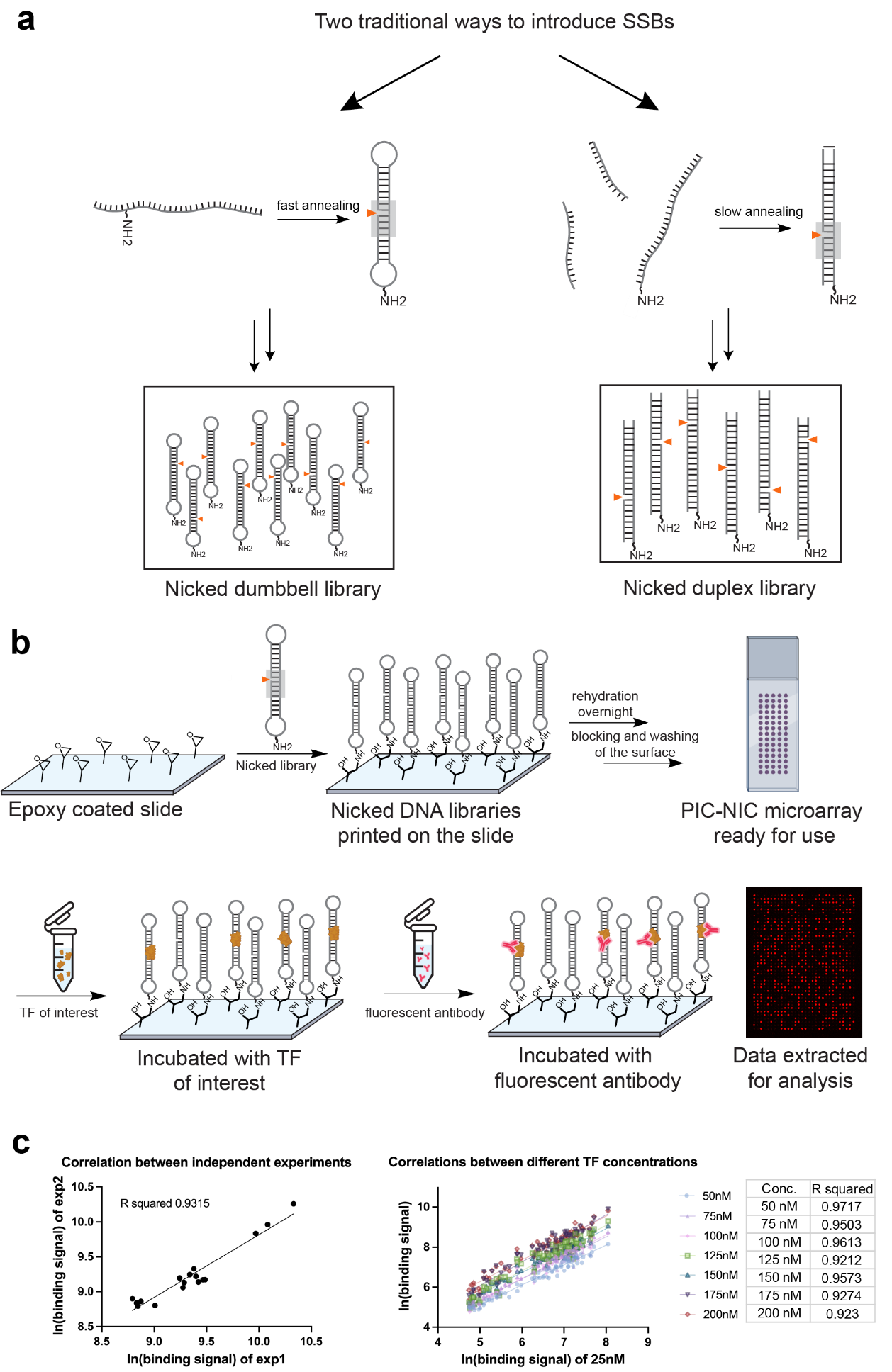

**Extended Data Fig. 1 PIC-NIC Pipeline and experimental reproducibility**

**a.** Two strategies to introduce SSBs into DNA TF binding sites using traditional approaches. *Left:* dumbbell-shaped DNA oligos fold on themselves upon fast annealing leaving a nick at the desired position; *Right:* slow annealing of three single-stranded DNA oligos leaves a nick at the desired position. Both methods were employed to yield oligo libraries of nicked TF binding sites. **b.** Schematic representation of the PIC-NIC experimental pipeline, for further details see Supplementary Methods. **c.** PIC-NIC experiments are highly reproducible and robust. *Left:* Binding measurements for the XYZ protein show strong correlation between two independently synthesized replicate microarrays (Pearson’s R² = 0.9315). *Right*: Data acquired at different TF concentrations remain highly consistent, demonstrating robustness across binding regimes.

**Extended Data Fig. 2 Structural families and contact maps of the 15 TFs employed in this study**

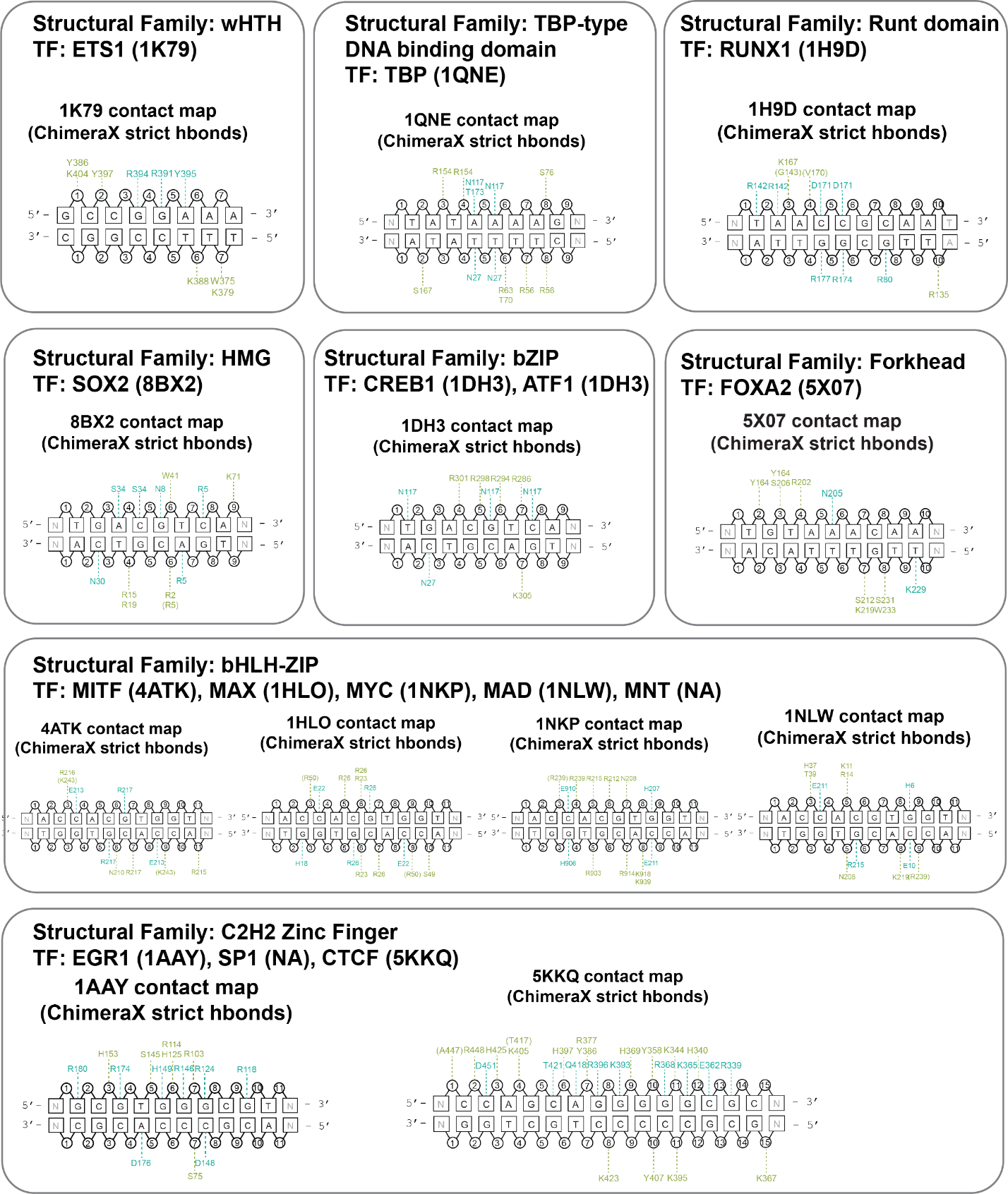

**Extended Data Fig. 2 Structural families and contact maps of the 15 TFs employed in this study.**

Structural families, PDB ID and detailed protein-DNA H-bond contact maps for the 15 TFs showing backbone contacts (green) and base contacts (turquoise). Contact maps were produced in ChimeraX employing a general cutoff distance of 3.5 Å (tolerance 0.4 Å) and angle cutoff of 100° (tolerance 20°). The binding site shown in the figure is the same length as the binding site chosen for PIC-NIC experiments.

**Extended Data Fig. 3 PIC-NIC binding profiles for the 15 TFs.**

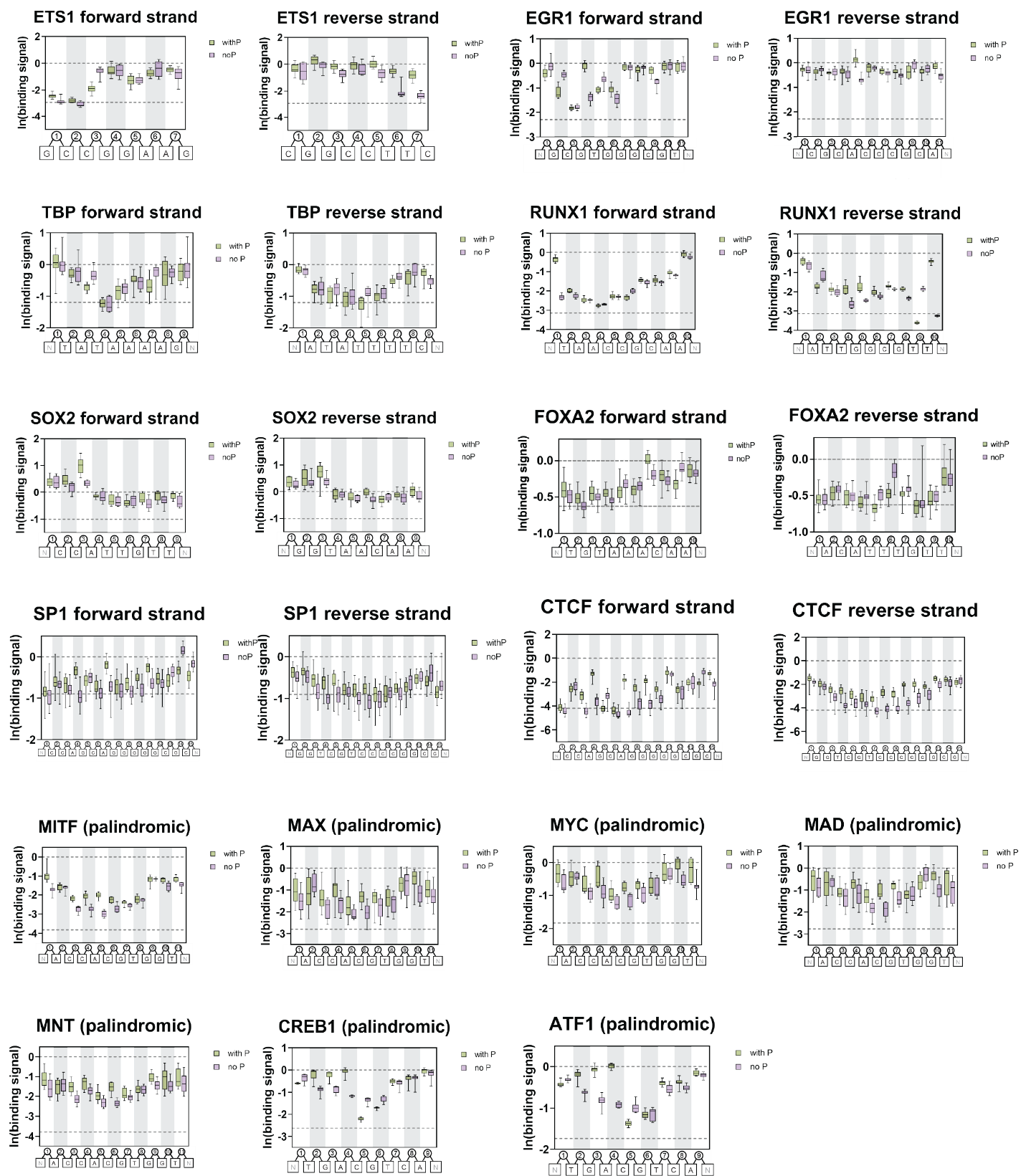

**Extended Data Fig. 3 PIC-NIC binding profiles for the 15 TFs.**

Box-and-whisker plots of binding signal of each TF to its respective binding site containing a nick at the indicated position, either with retention of 5’ phosphate (green) or removal of the phosphate (purple). The upper dashed line (y=0) represents the binding signal of the target TF to the intact DNA binding site. The lower dashed line represents the binding signal of the target TF to a non-specific site. Each box represents 10-20 spatially scattered replicates of the same DNA binding site on the array.

**Extended Data Fig. 4 Bio-Layer Interferometry (BLI) data for ETS1 and SOX2**

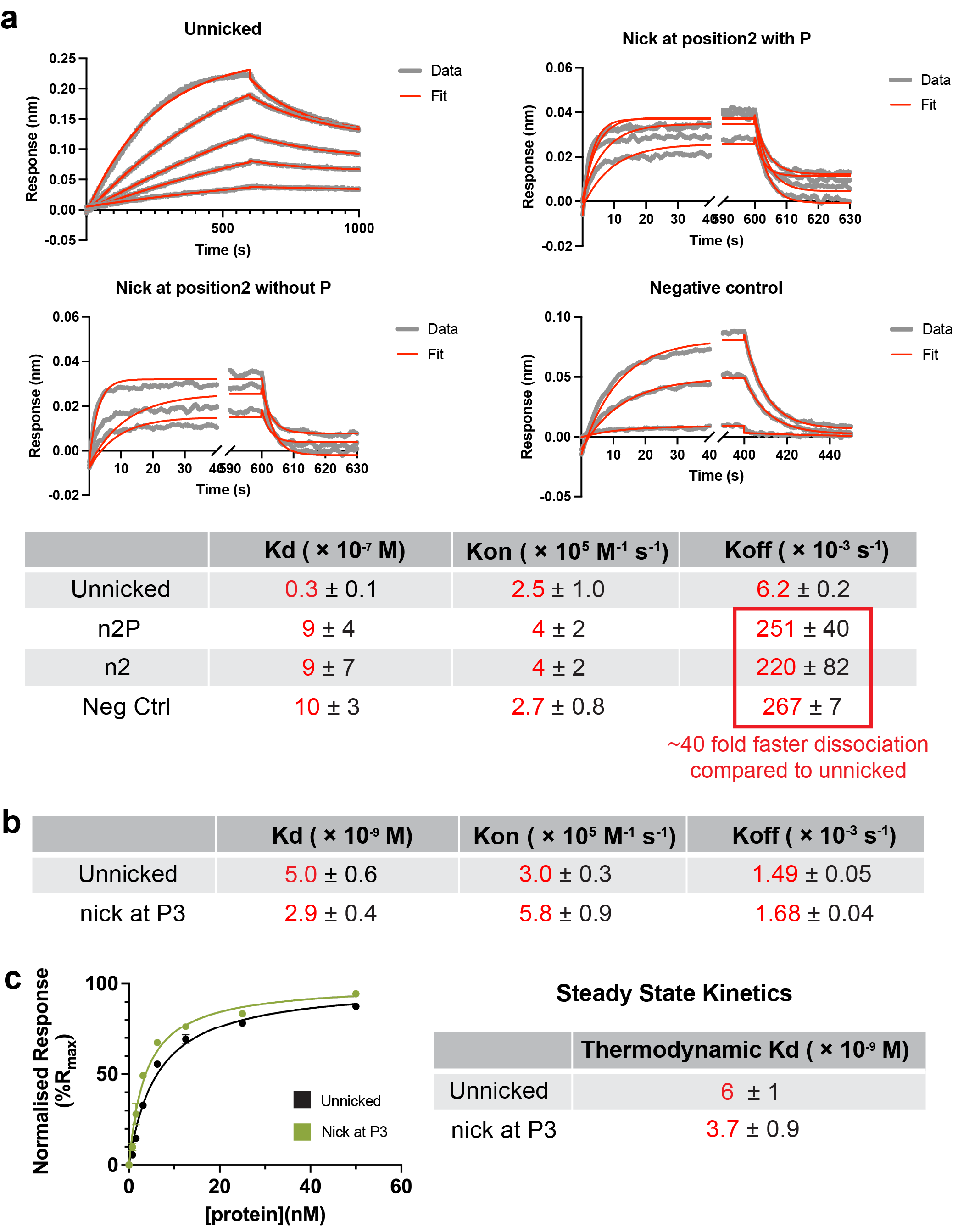

**Extended Data Fig. 4 Bio-Layer Interferometry (BLI) data for ETS1 and SOX2**

**a.** Bio-Layer Interferometry (BLI) sensorgrams (grey) of WT ETS1 binding to the intact (unnicked) DNA probe, the DNA probe nicked at position 2 (n2P), the DNA probe nicked at position 2 with the removal of 5’ phosphate (n2), and a probe containing a non-specific site (neg ctrl). Sensorgrams at each concentration were fit locally to a 1:1 binding model (red) at each concentration to determine *k*_on_ and *k*_off_ values. *K*_D_ values were calculated for each curve, and the means and standard deviations derived from these values are presented. Compared to the intact DNA probe, the binding to n2P, n2 and neg ctrl exhibited ~40-fold faster dissociation. **b.** Kinetic *K*_D_, *k*_on_ and *k*_off_ values derived from sensorgrams of SOX2 binding to the intact DNA probe and DNA probe nicked at position 3 (Fig. 4c). *k*_on_ and *k*_off_ values were obtained by global fitting of sensorgrams to a 1:1 binding model. **c.** Thermodynamic *K*_D_ of SOX2 binding to the intact DNA probe and DNA probe nicked at position 3 were calculated from 7 concentrations using a one-site binding model at equilibrium. The observed ~1.6-fold change in the thermodynamic Kd aligns with the ~1.7-fold change in the obtained kinetic *K*_D_ above.

**Extended Data Fig. 5 All atom Molecular Dynamic (MD) simulations for ETS1**

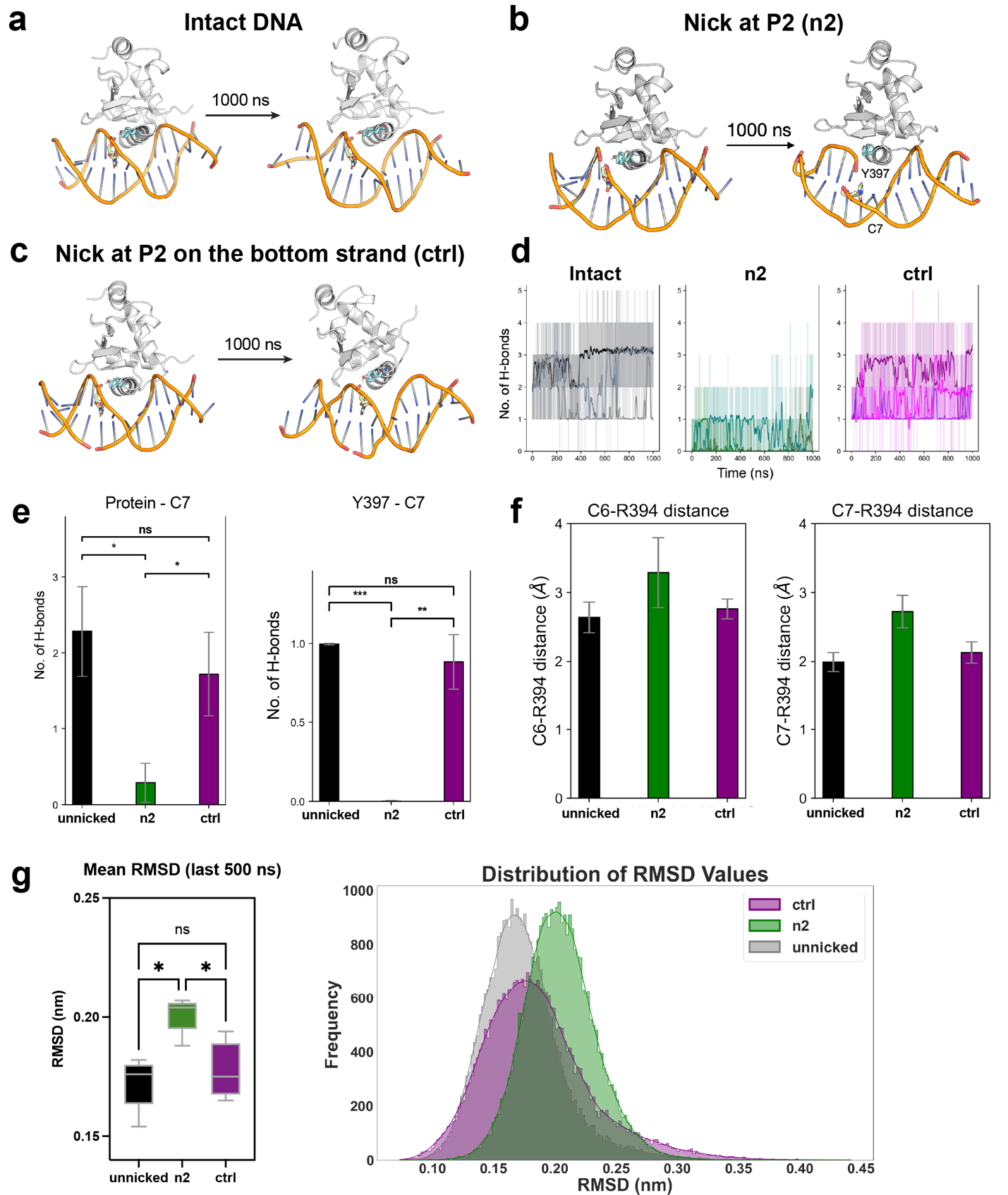

**Extended Data Fig. 5 All atom Molecular Dynamic (MD) simulations for ETS1**

**a.** MD simulation snapshots of wild-type ETS1 bound to intact DNA over a 1000 ns trajectory, serving as a structural reference for nick-induced perturbation analysis. **b.** MD simulation snapshots of wild-type ETS1 bound to DNA nicked at position 2 (with the removal of 5’ phosphate), where the nick abolishes binding, over a 1000 ns trajectory. Distinctive local distortion of the DNA backbone geometry is observed at the break site. **c.** MD simulation snapshots of wild-type ETS1 bound to DNA nicked at position 2 at the bottom strand (with the removal of 5’ phosphate), where the nick does not induce a major effect in binding of the TF, over a 1000 ns trajectory, serving as negative control. **d.** Time-dependent H-bond analysis across the simulation trajectory for the three constructs. In n2, the removal of the phosphate in contact with Y397 leads to the overall loss of hydrogen bonds over the 1000 ns trajectory. **e.** Hydrogen bonds formed between the protein and DNA at C7 (left), and specifically between Y397 and C7 (right), across the simulation trajectories. The n2 nick significantly reduces hydrogen bonding, unlike the control nick, and this effect is caused by the removal of C7 backbone phosphate at the break site. **f.** Average distances from C6 and C7 to R394 reveal that the n2 nick increases these distances relative to the wild-type and control, weakening interactions and reflecting geometric disruption of the protein–DNA interface. **g.** *Left panel*: Root-mean-square deviation (RMSD) of backbone atoms in the last 500 ns of simulation (assuming equilibrium has been reached) shows that the n2 nick increases structural fluctuations compared to the wild-type and control. *Right panel*: Distribution of RMSD values for the last 500 ns trajectory shows a rightward shift in the n2 nicked complex (green), indicating elevated structural flexibility compared to the wild-type (gray) and control nick (magenta).

**Extended Data Fig. 6 ETS1 mutants at phosphate-contacting residues exhibit differential binding in comparison to the WT protein**

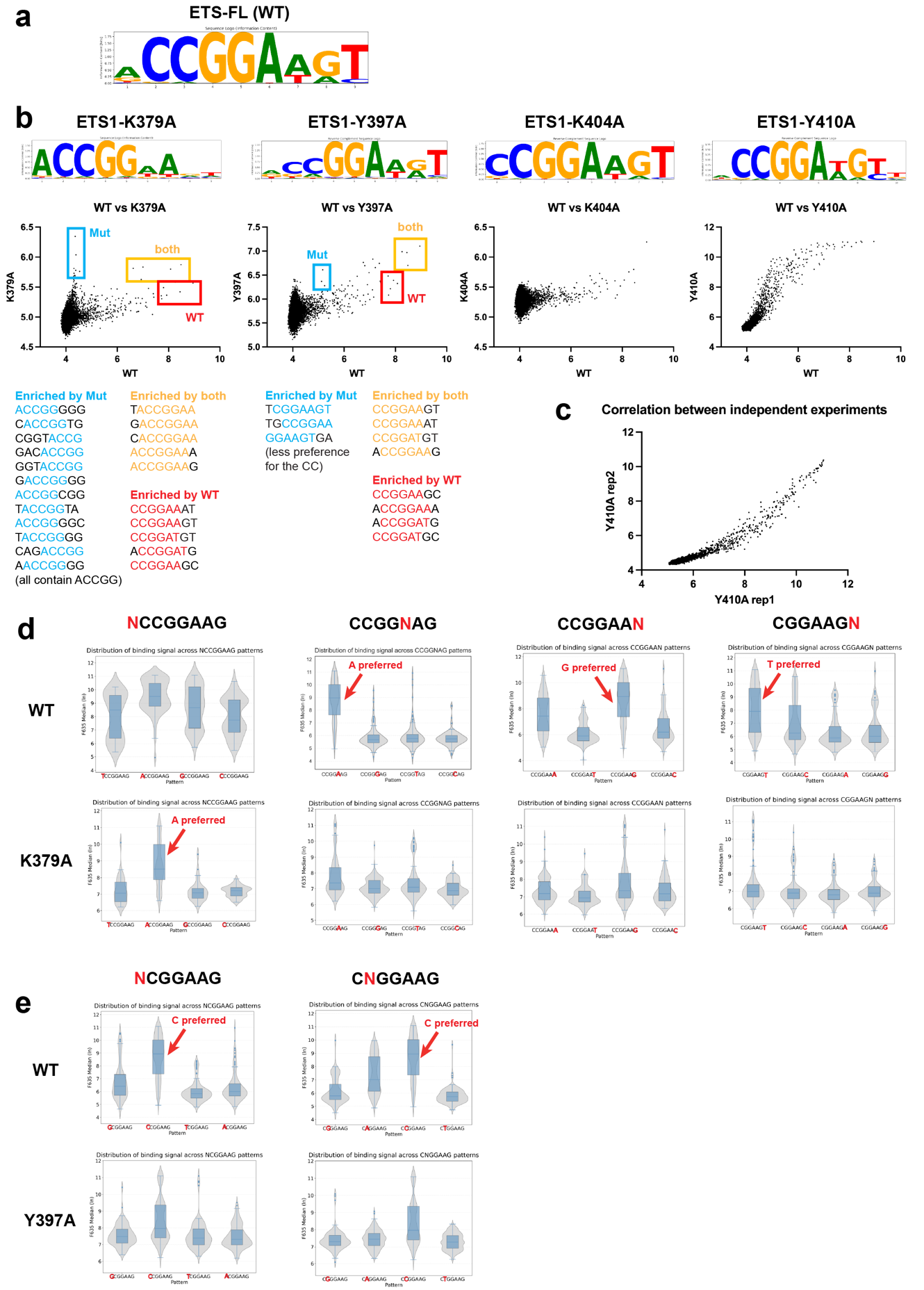

**Extended Data Fig. 6 ETS1 mutants at phosphate-contacting residues exhibit differential binding in comparison to the WT protein**

**a.** PWM logo of full-length wild-type ETS1 protein, generated from experimental data of the universal protein binding microarray containing all possible DNA 10-mer sequences. **b.** *Top panels*: PWM logos of four ETS1 mutants (K379A, Y397A, K404A and Y410A), generated using an identical universal 10-mer microarray design. *Middle panels*: Binding signal correlations of the four ETS1 mutants versus the ETS1 WT. Each point represents a DNA 8-mer sequence. Blue box highlights the 8-mer sequences preferred by the respective ETS1 mutant; red box highlights the 8-mer sequences preferred by the WT ETS1; yellow box highlights the 8-mer sequences highly bound by both the mutant and WT ETS1. *Bottom panels*: 8-mer sequences corresponding to the highlighted boxes in the middle panels. **c.** Correlation of replicate experiments (from independent protein expression) confirms high reproducibility. **d-e.** Comparison of nucleotide preference at this indicated position (N) for WT ETS1 and **d** K379A and **e** Y397A**.**

**Extended Data Fig. 7 High-resolution crystal structure analysis of TBP in complex with nicked DNA**

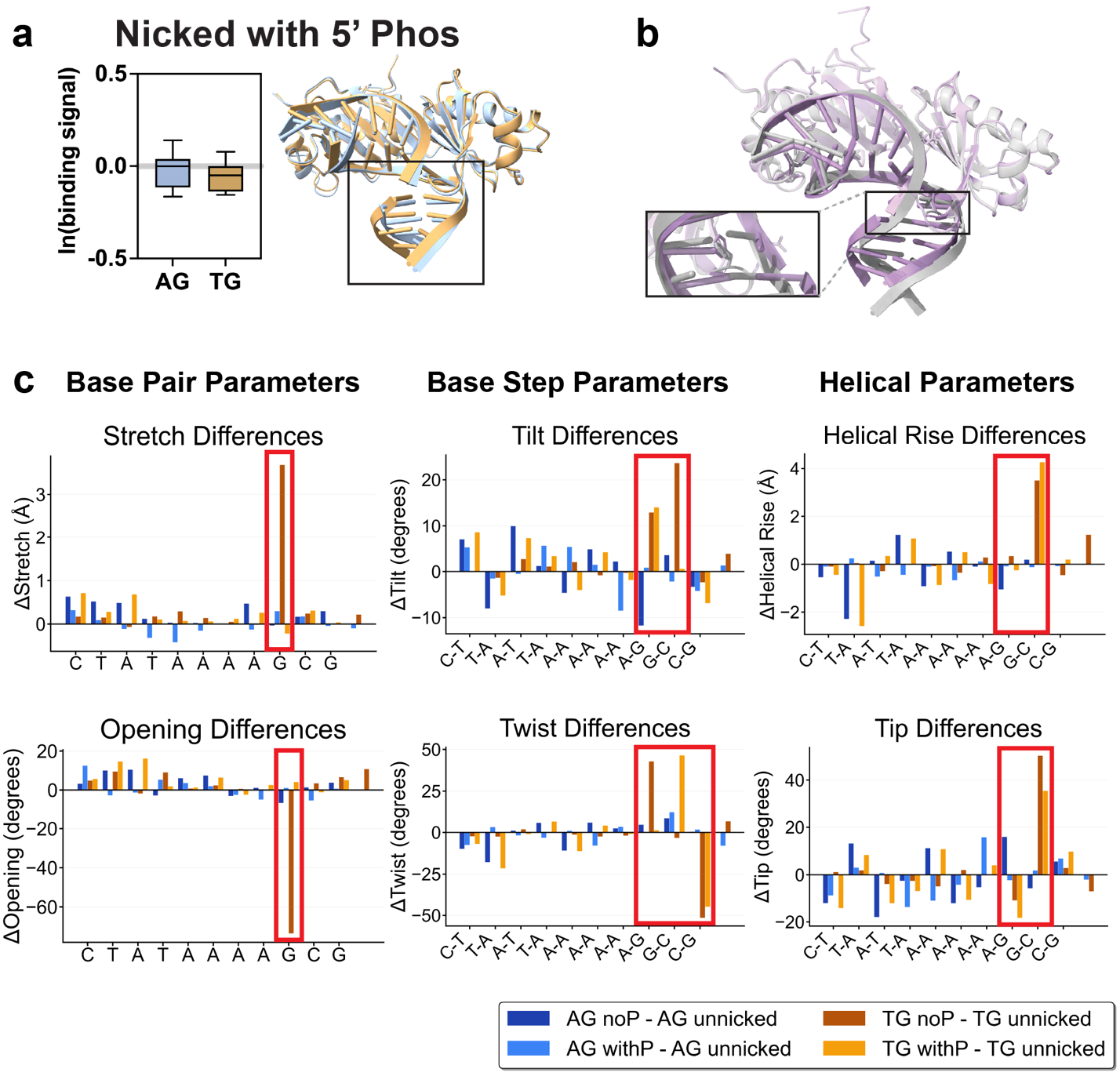

**Extended Data Fig. 7 High-resolution crystal structure analysis of TBP in complex with nicked DNA**

**a.** The binding level of TBP to the AG variant becomes almost identical to that of the TG variant when position 8 at the bottom strand is nicked with the retention of 5’ phosphate. Two high-resolution crystal structures of nicked complexes corresponding to the AG variant and the TG variant were solved (Blue: AG variant complex; gold: TG variant complex). Structural overlay reveals a larger DNA bending angle in the TG complex compared to the same region in the AG complex. **b.** Structural overlay and zoom in of the TG nicked complex (with the removal of 5’ phosphate at the break site) and TG intact complex (gray: intact TG complex; purple: nicked TG complex with no 5’ phosphate). When nicked, the DNA geometry deviates significantly from that of intact DNA, introducing a larger bending angle and more deformation. The structural zoom in shows dramatic local shifts in the base pair geometry flanking the nick site, suggesting that the nick disrupts the local DNA architecture. **c.** Structural analysis of the four solved crystal structures shows that the TG nicked complex with no 5’ phosphate has the largest deformation at the nick site. Representative base pair parameters (stretch and opening), base step parameters (tilt and twist) and helical parameters (helical rise and tip) were chosen to illustrate the parameter difference between nicked and intact complexes.

**Extended Data Fig. 8 High-resolution crystal structure analysis of EGR1 in complex with nicked DNA**
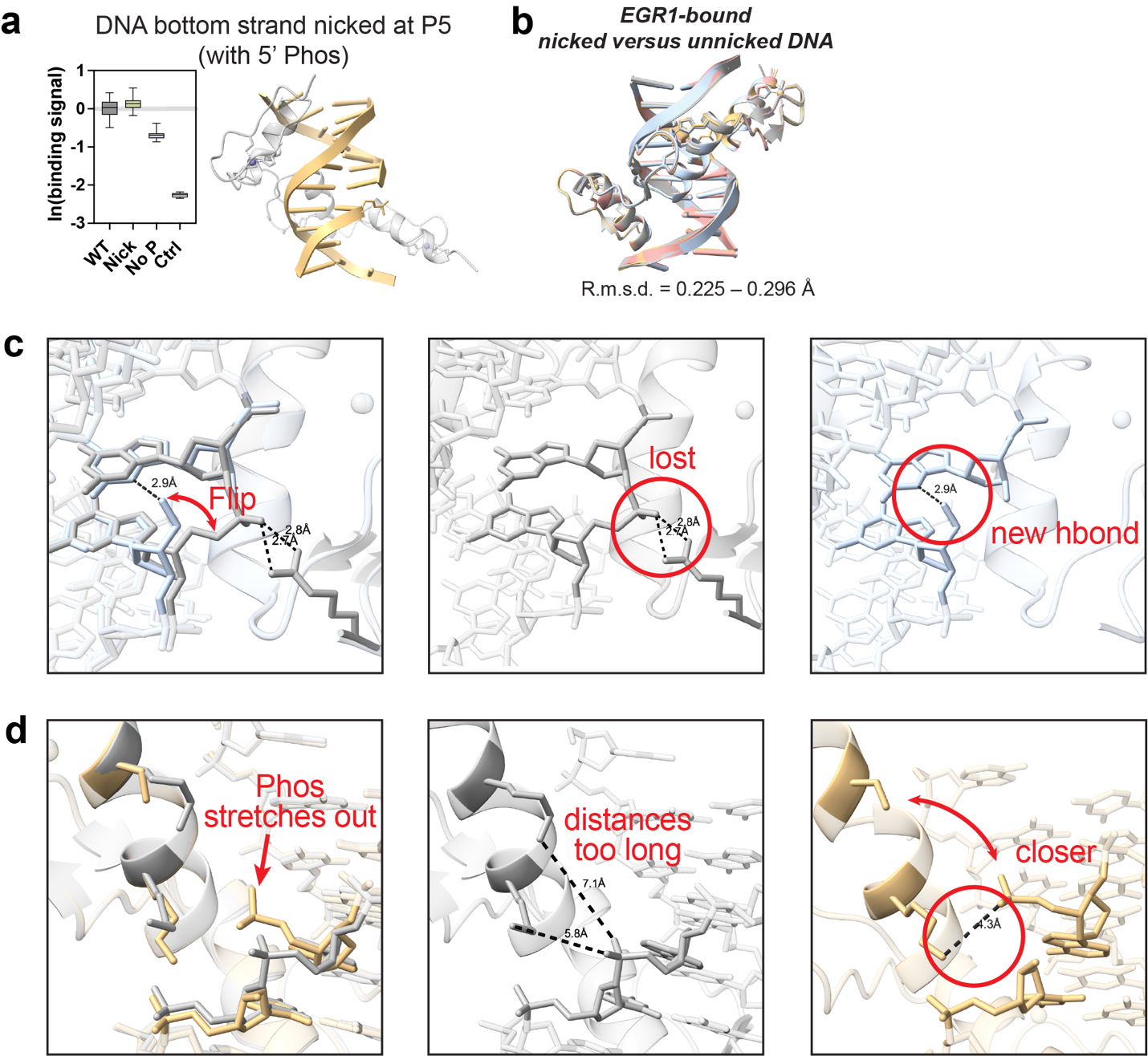

**Extended Data Fig. 8 High-resolution crystal structure analysis of EGR1 in complex with nicked DNA**

**a.** No disruption of EGR1 binding is observed when the DNA is nicked at position 5 on the bottom strand (5′ phosphate retained), while when the 5’ phosphate is removed, only minor disruption is observed. The high-resolution crystal structure of EGR1 in complex with DNA nicked at position 5 on the bottom strand (5’ phosphate retained) is solved. **b.** Overlay of EGR1 bound to intact and three nicked DNA duplexes reveals high structural similarity (RMSD = 0.225 - 0.296 Å), indicating minimal global deformation. **c.** Structural changes when the DNA is nicked at position 7 on the top strand (phosphate removed). Gray: EGR1/intact DNA complex; blue: EGR1/nicked DNA complex. *Left panel*: Compared to the intact DNA, the backbone at the break site in the nicked DNA flips direction and induces a conformational change. *Middle panel*: The original H-bonds between Arg103 and the backbone phosphate are lost due to the phosphate removal at this position. *Right panel*: Due to the flipping of the backbone in the nicked structure, a new intramolecular H-bond is formed between the backbone OH and the neighboring guanine base. **d.** Structural changes when the DNA is nicked at position 5 on the bottom strand (phosphate retained). Gray: EGR1/intact DNA complex; gold: EGR1/nicked DNA complex. *Left panel*: When nicked, the backbone phosphate stretches away from its original position and the backbone geometry around the nicked phosphate is altered. *Middle panel*: In the intact DNA complex, the distances between the amino acid residues and DNA phosphate backbone at this position are too long to contribute to stabilizing interactions. *Right panel*: Due to the conformational stretch of the phosphate backbone in the nicked complex, the distances between nearby Arg and Lys residues and this backbone phosphate are shortened, and thus new favorable interactions emerge.

**Extended Data Table. 1 Crystallographic data collection and refinement statistics: TBP-**

**DNA complexes**

|  | **TBP-TG-P** | **TBP-TG** | **TBP-AG-P** | **TBP-AG** |
| --- | --- | --- | --- | --- |
| **Data collection** |  |  |  |  |
| PDB code  Space group  Wavelength (Å)  Resolution (Å) | 9OW8  P2_1_  1.01  48.6-2.60  (2.69-2.60)* | 9OW7  C222_1_  1.01  65.7-2.30  (2.45-2.30) | 9OWZ  I2  1.01  88.3-2.91  (3.10-2.91) | 9OWI  P2_1_  1.01  48.3-2.49  (2.52-2.48) |
| Cell dimensions |  |  |  |  |
| *a*, *b*, *c* (Å) | 69.1,117.5,102.1 | 77.6,123.4,66.4 | 122.7,106.2,161.9 | 68.6,117.3,102.1 |
| α, β, γ (°) | 90.0,106.9,90.0 | 90.0,90.0,90.0 | 90.0,101.1,90.0 | 90.0,107.0,90.0 |
| R-merge  R-pim | 0.083 (0.508)  0.055 (0.321) | 0.053 (0.644)  0.027 (0.350) | 0.076 (0.595)  0.049 (0.384) | 0.086 (0.550)  0.055 (0.342) |
| Mean I/sigma (I) | 9.6 (2.3) | 13.6 (1.4) | 8.3 (1.9) | 9.0 (2.2) |
| Completeness (%) | 97.7 (99.5) | 98.6 (90.5) | 97.5 (98.4) | 99.1 (98.3) |
| Redundancy  CC_1/2_ | 3.3 (3.4)  0.994 (0.794) | 5.8 (5.3)  0.998 (0.851) | 3.3 (3.4)  0.994 (0.679) | 3.5 (3.5)  0.995 (0.759) |
| **Refinement** |  |  |  |  |
| No. reflections | 46965 (4418) | 17896 (328) | 43821 (1466) | 54270 (4434) |
| *R*_work_ / *R*_free_ (%) | 20.2/25.3 | 18.9/22.9 | 23.5/25.3 | 20.7/25.1 |
| R.m.s. deviations |  |  |  |  |
| Bond lengths (Å) | 0.009 | 0.008 | 0.012 | 0.008 |
| Bond angles (°)  Ramachandran  Favored (%)  Disallowed (%)  Clashscore | 1.05  96.0  0.00  18.2 | 1.00  99.0  0.00  9.69 | 1.28  91.6  0.00  21.93 | 0.965  95.9  0.00  14.0 |

*Values in parentheses are for the highest-resolution shell.

**Extended Data Table. 2 Crystallographic data collection and refinement statistics: EGR1-**

**DNA complexes**

|  | **EGR1-n7** | **EGR1-ren5-P** | **EGR1-ren7** |
| --- | --- | --- | --- |
| **Data collection** |  |  |  |
| PDB code  Space group  Wavelength (Å) | 9RIC  C222_1_  1.34 | 9RI6  C222_1_  1.34 | 9RJ6  C222_1_  1.34 |
| Resolution (Å) | 17.23-1.9  (2.09-1.90) | 17.43-2.0  (2.07-2.00) | 18.38-1.95  (2.15-1.95) |
| Cell dimensions  *a*, *b*, *c* (Å)  α, β, γ (°) | 44.56,56.05,130.21  90,90,90 | 44.77,55.51,129.88  90,90,90 | 44.45,56.16,129.79  90,90,90 |
| R-merge | 0.053 (0.207) | 0.073 (0.237) | 0.100 (0.458) |
| R-pim | 0.021 (0.085) | 0.028 (0.098) | 0.037 (0.169) |
| Mean I/sigma (I) | 23.3 (7.5) | 19.11 (6.5) | 14.33 (5.2) |
| Completeness (%) | 99.82 (99.95) | 99.88 (100.00) | 99.85 (100.00) |
| Redundancy  CC_1/2_ | 7.4 (7.0)  0.999 (0.977) | 7.2 (6.7)  0.998 (0.953) | 8.4 (7.8)  0.997 (0.945) |
| **Refinement** |  |  |  |
| No. reflections | 13,206 (3,254) | 11,283 (2,773) | 12,198 (2,997) |
| *R*_work_ / *R*_free_ (%) | 19.2/24.1 | 18.6/21.7 | 19.9/21.2 |
| R.m.s. deviations  Bond lengths (Å) | 0.007 | 0.008 | 0.007 |
| Bond angles (°) | 0.89 | 0.93 | 0.91 |
| Ramachandran  Favored (%)  Disallowed (%) | 100.00  0.00 | 98.8  0.00 | 100.00  0.00 |
| Clashscore | 0.48 | 1.44 | 0.48 |
