## Supplementary Information for "Systematic DNA nicking reveals the structural logic of protein recognition"

Singapore;

^5^Department of Biochemistry, Duke University School of Medicine, Durham, NC 27710, USA

**This PDF file includes:**

Supplementary Methods

Titles and Legends for Supplementary Tables 1-8

Supplementary References

**Supplementary Methods**

**Protein expression and purification**

Protein expression and purification for PIC-NIC assays:

Human ETS1 full-length protein was expressed as an N-terminal GST fusion and purified in *E. coli*^1^. Bacterial cultures were grown at 37 °C in LB media, induced overnight at 15 °C with 0.5 mM IPTG, harvested, and lysed by sonication in PBS. The clarified lysate was applied to a 5 mL GSTrap FF column (Cytiva) equilibrated in 25 mM Tris-HCl pH 8, 50 mM KCl, 10% (v/v) glycerol, 0.1 mM EDTA, and 1 mM DTT. The protein was eluted in 50 mM Tris-HCl (pH 8.0), 20 mM KCl, 20 mM reduced glutathione. Further purification was achieved by heparin chromatography (HiTrap Heparin HP, Cytiva), eluting over a linear gradient from 50 mM Tris-HCl (pH 8.0) to 50 mM Tris-HCl (pH 8.0), 1 M KCl, with GST-ETS1 eluting at ~300 mM KCl. The protein was then subjected to size-exclusion chromatography (Superdex 200 10/300 GL, Cytiva) equilibrated in PBS. Protein-containing fractions were pooled, and the pure proteins were concentrated to 0.0051 mM (0.399 mg/mL). The concentration was determined by UV absorption at 280 nm using an extinction coefficient of 124525 M^-1^ cm^-1^. The protein was snap frozen in liquid N_2_ and stored at -80 °C.

Human EGR1, residues 335-423, with N-terminus GST tag, were expressed and purified as described below in EGR1 expression and purification for crystallography, without thrombin cleavage.

*Arabidopsis thaliana* TBP, with N-terminal HIS tags, were expressed and purified in *E. coli* C41 (DE3) cells as described previously^2^. In brief, an overnight culture was inoculated into LB media and grown at 37 °C. Expression was induced by 0.5 mM isopropyl β-D-1-thiogalactopyranoside overnight at 15 °C. Cells were harvested and resuspended in a buffer containing 5% (v/v) glycerol, 4 mM MgCl_2_, 600 mM NaCl and 40 mM MES (pH 7.2). Cells were lysed by French press and sonication followed by centrifugation. The lysate was purified on a Ni-NTA column via the N-terminal 6x HIS tag, washed with the lysis buffer with increasing concentration of imidazole. TBP was then eluted in the lysis buffer with 100-1000 mM imidazole, and 5 mM BME was added to the fractions. The eluent was dialysed against a buffer containing 20 mM HEPES-KOH (pH 8), 100 mM KCl, 20% (v/v) glycerol, 1 mM MgCl_2_ and 1 mM CaCl_2_. The protein was concentrated with a 10kDa MWCO Amicon (Millipore) and the concentration was determined by UV absorption at 280 nm using an extinction coefficient of 10.5 mM^-1^ cm^-1^. The protein was snap frozen in liquid N_2_ and stored at -80 °C.

Full length human Max, DNA-binding doman of Myc, Mad, and Mnt were chemically synthesised and labelled with TAMRA fluorophore, as described previously^3,4^. They were generously provided by Dr. Muhammad Jbara (Tel Aviv University). All experiments of Myc, Mad, and Mnt heterodimers were performed by mixing Max with its desired binding partner, at a 1:5 ratio, to ensure that mostly heterodimers were formed instead of Max:Max homodimers.

Recombinant human full-length Runx1 protein was expressed and purified from BL21 (DE3) *E. coli* harboring a clone in pReceiver-B03 Runx1. In brief, an overnight culture was inoculated into LB media and grown at 33 °C until OD reached 0.6-0.8. Expression was induced by 0.5 mM isopropyl β-D-1-thiogalactopyranoside overnight at 15 °C. Cells were harvested and resuspended in a buffer containing 500 mM Tris (pH 7.5), 300 mM NaCl, 1 mM DTT and 5 mM imidazole. Cells were lysed by French press and sonication followed by centrifugation. The lysate was purified on Ni-NTA column via the N-terminal 6x HIS tag, washed with a buffer containing 50 mM Tris (pH 7.5), 300 mM NaCl, 5 mM 2-Mercaptoethanol (BME), 5 mM imidazole and 10% (v/v) glycerol. The protein was eluted with an elution buffer consisting of 50 mM Tris (pH 7.5), 300 mM NaCl, 5 mM 2-Mercaptoethanol (BME), 200 mM imidazole and 10% (v/v) glycerol. The eluate was dialysed against the elution buffer without imidazole overnight, followed by a heparin Sepharose SP (GE) column. The protein was then diluted with a buffer consisting of 50 mM Tris (pH 7.5), 2-Mercaptoethanol (BME) and 10% (v/v) glycerol to 100 mM salt concentration, loaded onto a heparin Sepharose SP (GE) column and washed with the same buffer. The protein was then eluted with a salt gradient to a buffer consisting of 1 M NaCl, 50 mM Tris (pH 7.5), 2-Mercaptoethanol (BME) and 10% (v/v) glycerol.  The eluate was concentrated and the concentration was determined by UV absorption at 280 nm using an extinction coefficient of 40340 M^-1^ cm^-1^. The protein was snap frozen in liquid N_2_ and stored at -80 °C.

Recombinant human Sox2 DNA binding domain, residues 19-118, with N-terminal HIS tags, was expressed and purified as described below in Sox2 expression and purification for BLI.

Human FoxA2 DNA-binding domain, residues 142-269, with N-terminal GST tags and C-terminal HIS tags, was expressed by *in vitro* transcription/translation system with PURExpress® *In Vitro* Protein Synthesis Kit (New England Biolabs, catalog #: E6800). A standard protocol was followed, with the addition of 10 μl of Solution A, 7.5 μl of Solution B, 1 μl of RNase Inhibitor and 250 ng template plasmid (per 25 μl reaction). The resulting reaction mixture was incubated at 37°C for 2 hours and stored at -20 °C.

Full-length human MITF, with N-terminal GST tag and C-terminal HIS tag, was purchased from Origene (catalog #: TP761396). Full-length human CREB1, with N-terminal HIS tag, was purchased from Origene (catalog #: TP760318). Human ATF1, residues 1-271, with N-terminal HIS tag, was purchased from Prospec (catalog #: PKA-019). Full-length human SP1 with N-terminal HIS tag was purchased from Origene (catalog #: TP760592). Full-length human CTCF with N-terminal GST tag was purchased from Abnova (catalog #: H00010664-P01).

ETS1 expression and purification for BLI: Recombinant ETS1 DNA-binding domain (murine residues 331 to 440) was expressed and purified from BL21*(DE3) *E. coli* harboring a clone in pET28b-Ets1-ETS (Addgene #85735) as previously reported^5^. In brief, an overnight culture was inoculated into liter-scale LB media and grown at 37 °C. Expression was induced by 1 mM isopropyl β-D-1-thiogalactopyranoside overnight at 25 °C. Cells were harvested and resuspended in a buffer containing 100 mM Tris-HCl (pH 8.0), 500 mM NaCl and 5 mM DTT. All downstream buffers contained 0.5 mM Tris(2- carboxyethyl)phosphine (TCEP) hydrochloride to reduce the cystein disulfide bridges. Cells were lysed by French press and sonication followed by centrifugation. 40 μL of IGEPAL CA-630 (Sigma) and DNA inhibitor were added to 40 mL of the lysate. The lysate was purified on Ni-NTA column via the C terminal 6xHis tag on the protein, followed by a heparin Sepharose SP (GE) column. The eluate was dialysed overnight against a buffer containing PBS, 0.15 M NaCl and bovine thrombin (Prospec) to remove the C-terminal 6×His tag. Purified Ets-1 was dialysed extensively into final buffer of 10 mM NaH_2_PO_4_/ Na_2_HPO_4_ (pH 7.4) with 0.15 M NaCl. Ets-1 concentration was determined by UV absorption at 280 nm.

Sox2 expression and purification for BLI: Recombinant human Sox2 DNA binding domain, residues 19-118, with N-terminal HIS tags, was expressed and purified from BL21 (DE3) *E. coli* as previously described^6^. In brief, an overnight culture was inoculated into LB media and grown at 37 °C. Expression was induced by 0.5 mM isopropyl β-D-1-thiogalactopyranoside for 3-4 hours at 37 °C. Cells were harvested and resuspended in a buffer containing 50 mM Tris-HCl (pH 7.5), 500 mM NaCl, 10 mM imidazole, 10% (v/v) glycerol and 5 mM 2-Mercaptoethanol (BME). Half a pill of cOmplete Protease Inhibitor (Sigma) and 1 mM DTT was added to the sample, before the cells were lysed by French press and sonication, followed by centrifugation at 14000 rpm for 30 min at 4 °C. A HisTrap™ High Performance (Cytiva) column was equilibriated with a buffer containing 50 mM Tris-HCl (pH 7.5), 500 mM NaCl, 10 mM imidazole, 10% (v/v) glycerol and 5 mM 2-Mercaptoethanol (BME). The supernatant was syringe filtered through a 0.45 μm filter, before the lysate was loaded onto the column at 1-1.5 ml/min. The column was wash with the same equilibriation buffer at 2-3 ml/min with several column volumes until the absorbance reading was close to the baseline. The protein was then eluted with a salt gradient to a buffer consisting of 50 mM Tris-HCl (pH 7.5), 500 mM NaCl, 500 mM imidazole, 10% (v/v) glycerol and 5 mM 2-Mercaptoethanol (BME), collected in fractions. Protein-containing fractions were pooled and the pure proteins were dialysed against a buffer consisting of 50 mM Tris-HCl (pH 7.5), 500 mM NaCl, 10% glycerol and 5mM 2-Mercaptoethanol (BME) overnight. The protein was concentrated and the concentration was determined by UV absorption at 280 nm using an extinction coefficient of 13980 M^-1^ cm^-1^. The protein was snap frozen in liquid N_2_ and stored at -80 °C.

TBP expression and purification for crystallography: For expression and purification of the *Arabidopsis thaliana* TATA box binding protein (TBP) protein, *E. coli* C41(DE3) cells were transformed with a pET15b plasmid encoding the *A. thaliana* TBP (generated as a codon-optimized gene for *E. coli* expression) with a cleavable hexahistidine-tag on the N-terminus (Genscript)^2^. For protein expression, cells were grown at 37 ºC to an OD_600_ of 0.5, induced at 15 ºC with 0.5 mM ITPG overnight and pelleted the next day. The cell pellets were reconstituted with buffer A (25 mM Tris pH 7.5, 500 mM NaCl, 5% (v/v) glycerol, 0.5 mM β-mercaptoethanol (BME)), lysed twice using a microfluidizer and pelleted. The supernatant was loaded onto a Cobalt-NTA column and the column was washed overnight with buffer A and then eluted with increasing concentrations of imidazole in buffer A. TBP eluted in buffer A containing between 100 and 300 mM imidazole. The resultant TBP was >95% pure at this stage. Concentrations were determined by UVvis using a calculated molar absorption coefficient of 10.5 mM^-1^ cm^-1^. For crystallization the hexahistidine tag was removed using a Thrombin cleavage capture kit. The his-tag free protein was concentrated just prior to crystal setups using a centricon 30 (30 MW cutoff).

EGR1 expression and purification for crystallography: Recombinant EGR1 DNA-binding domain (residues 335 to 423) was expressed and purified from BL21*(DE3) *E. coli* harboring a clone in pGEX 2T-egr1-DBD (Addgene #85735) as previously reported^7^. In brief, an overnight culture was inoculated into liter-scale LB media and grown at 37 °C until OD reached around 0.5. The temperature was then lowered to 22 °C, and after 20 min, expression was induced by the 1 mM isopropyl β-D-1-thiogalactopyranoside overnight at 22 °C with shaking (175 rpm). Cells were harvested by centrifugation at 4000 rpm at 4 °C for 20 min and resuspended in the lysis buffer containing 20 mM Tris (pH 7.5), 500 mM NaCl, 5% (v/v) glycerol, 0.5 mM TCEP, 25 μM ZnCl_2_ and PIC. Cells were lysed by French press and sonication, followed by centrifugation at 50,000 rpm at 4 °C for 30 min. 4% w/v solution (10X) poly(ethylenimine) in water, pH 8 was prepared by dissolving 1.6 g PEI (50% w/v) in 20 mL DDW. Solution was acidified to pH 8 by conc. HCl. 1/10 volume of 4% PEI solution was added to the supernatant, inducing the formation of a precipitate. The mixture was pelleted by centrifugation at 18,000 rpm at 4 °C for 10 min. GSTrap HP 5 mL column was pre-equilibrated in the lysis buffer above. The cleared extract sample was loaded onto the column and non-GST tagged material was eluted in 100% lysis buffer. The GST fusion proteins were then eluted with 20 mM glutathione (GSH) in the elution buffer containing 100 mM Tris-HCl (pH 8.0), 5% (v/v) glycerol, 25 μM ZnCl_2_, and 250 mM NaCl. The GST tag was removed using human Thrombin (Prospec, catalog #: PRO-339), incubated at room temperature with low-speed magnetic stirrer for 20 hours. The cleavage protein was diluted with 20 mM Tris (pH 7.5), 5% (v/v) glycerol, 25 μM ZnCl_2_, and 0.5 mM TCEP buffer to lower salt concentration. HiTrap Heparin HP 5 mL column was pre-equilibrated in 20 mM Tris (pH 7.5), 5% (v/v) glycerol, 25 μM ZnCl_2_, and 0.5 mM TCEP. The sample was loaded onto the column, eluted with a linear gradient to 1 M NaCl, collecting in fractions. The fractionated protein was pooled and concentrated with 3kDa MWCO Amicon (Millipore) at 4000 rpm at 4 °C to a volume of approximately 3 mL. Next, a Superdex 75 26/60 size exclusion column was pre-equilibrated overnight in 500 mM NaCl, 20 mM Tris-HCl (pH 7.5), 5% (v/v) glycerol, and 25 μM ZnCl_2_. The sample was loaded onto the column and eluted as a single peak in 500 mM NaCl, 20 mM Tris-HCl (pH 7.5), 5% (v/v) glycerol, and 25 μM ZnCl_2_. The eluate was concentrated and the concentration was determined by UV absorption at 280 nm using an extinction coefficient of at 1865 M^-1^ cm^-1^. The protein was snap frozen in liquid N_2_ and stored at -80 °C. Dynamic Light Scattering (DLS) measurement was carried out every time before the use of the protein to ensure there is no aggregation of the protein.

FOXA2 Plasmid Construction

The FOXA2 expression plasmid was constructed using the pET-29 backbone (Twist Bioscience). The insert included, in-frame and sequentially, an N-terminal GST tag, a TEV protease cleavage site, the human FOXA2 Forkhead domain (residues 142–269)^8^, a second TEV site, and a C-terminal His₆ tag. The construct was synthesized and cloned by Twist Bioscience. The final plasmid was sequence-verified prior to downstream applications.

**Site-directed mutagenesis**

The primers used to generate ETS1 mutants are as follows:

K379A: 5' - GGAGATGGGGAAAGAGGGCAAACAAACCTAAGATG - 3'

Y397A: 5' - GCCGTGGCCTACGCTACTATGCCGACAAAAACATCATCCAC - 3'

K404A: 5' - CGACAAAAACATCATCCACGCGACAGCGGGGAAACGCTACG - 3'

Y410A: 5' - GCGGGGAAACGCGCCGTGTACCGCTTTG - 3'

**DNA microarray production for PIC-NIC assays**

Sample Preparation

Homemade microarrays were produced in-house for PIC-NIC assays. Oligonucleotides were purchased from Integrated DNA Technologies (IDT) and W.M. Keck Biotechnology Resource Lab (Yale University). For nicks created from dumbbell-shaped DNA, single-stranded oligos were ordered with an internal amino modifier /iAmMC6T/, as well as a choice of 5' phosphate (/5Phos/). The sequences were designed to self-anneal into a dumbbell, leaving a nick at the desired position. The purchased oligos were each solubilised to 100 μM concentration according to the manufacturer datasheet, before accurate concentration quantification with Thermo Scientific Nanodrop One Microvolume UV-Vis Spectrometer. The oligos were next each diluted to a concentration of 5 μM in the spotting buffer consisting of 100 mM Na_2_CO_3_/NaHCO_3_ (pH 8.5). The oligo was then heated to 95 ºC for 2 min, followed by fast cooling on ice. The annealed oligos were centrifuged for 5 min, and 20 μl of the clear solution was plated in sciSOURCEPLATE 384 PS (Scienion). For nicked duplex oligos constructed from three DNA strands, the longest DNA part was ordered with an amino modifier at the end of the oligo (/5AmMC6/ or /3AmMC6T/). The purchased oligos were each solubilised to 100 μM concentration according to the manufacturer datasheet, before accurate concentration quantification with Thermo Scientific Nanodrop One Microvolume UV-Vis Spectrometer. The oligos were next mixed to a final construct concentration of 5 μM with the desired stoichiometry (1:5:5, with the long and labelled strand being 1, to ensure this strand is fully consumed) in the buffer consisting of 100 mM Na_2_CO_3_/NaHCO_3_ (pH 8.5). The oligo construct was then heated to 95 ºC for 2 min, followed by slow gradient cooling over 2 hours to ensure proper annealing. The annealed oligos were centrifuged for 5 min, and 20 μL of the clear solution was plated in sciSOURCEPLATE 384 PS (Scienion).

PIC-NIC DNA microarray production

The prepared DNA samples were printed onto the chips with our in-house sciFLEXARRAYER S12 automated non-contact dispensing system (Scienion). The empty slides used for printing were glass slides coated with epoxy surface (sciCHIP Epoxy from Scienion, and 3-D Epoxy from PolyAn). The layout of the microarray were designed, with 10-50 replicates of each material in the plate. The plated materials were spotted onto the glass slide at a temperature between 20-24 ºC and 70 ± 5 % relative humidity. Each droplet was dispensed at a volume of 300 ± 30 pL, forming a DNA spot on the glass slide with diameter of 90 ± 10 μm. After spotting, the DNA microarray was placed in a homemade humidity chamber filled with 3 M KCl (~70% humidity) overnight for rehydration and immobilisation. The DNA microarray was then placed in a coplin jar filled with PolyAn Blocking Buffer A for an hour, to deactivate the unreacted epoxy groups on the glass slide. Next, the microarray was subjected to a salt gradient wash in 5 coplin jars filled separately with PolyAn Washing Buffer I,  PolyAn Washing Buffer II, PolyAn Washing Buffer III, DDW and DDW. Each wash was 5 min, shaking at 125 rpm. The resulting microarray was spun dry in a slide spinner for 3 min (Labnet International, Inc.) and stored in a vacuum dessicator until further use.

**Protein binding and antibody steps for PBM and PIC-NIC assays**

Protein binding reactions were performed in the same conditions as previously described in PBM protocols^9,10^. The binding buffer, unless otherwise specified, contains PBS / 2% (w/v) milk / 51.3 ng/μl salmon testes DNA (Sigma) / 0.2 μg/μl bovine serum albumin (NEB) / 1 mM dithiothreitol. For EGR1, SP1, and CTCF, the binding buffer was: 10 mM Tris-HCl (pH 7.5), 150 mM KCl, and 0.2 μM ZnCl_2_, 2% (w/v) milk, 51.3 ng/μl salmon testes DNA, 0.2 μg/μl bovine serum albumin, 1 mM dithiothreitol. For TBP, the binding buffer was 10 mM HEPES, 70 mM KCl, 10 mM MgCl_2_, 1 mM EDTA, 2% (w/v) milk, 51.3 ng/μl salmon testes DNA, 0.2 μg/μl bovine serum albumin, 1 mM dithiothreitol. For CREB1, the binding buffer was 25 mM Tris-HCl (pH 7.4), 0.5 mM EDTA, 2% (v/v) glycerol, 5 mM MgCl_2_, 50 mM KCl, 25 mM boric acid, 2% (w/v) milk, 51.3 ng/μl salmon testes DNA, 0.2 μg/μl bovine serum albumin, 1mM dithiothreitol. For Sox2, the binding buffer was 10 mM HEPES (pH=7.5), 1 mM MgCl_2_, 0.01 mM ZnCl_2_, 10 mM NaCl, 2% (wt/vol) milk, 51.3 ng/μl salmon testes DNA, 0.2 μg/μl bovine serum albumin, 5% glycerol, 1 mM dithiothreitol. For FoxA2, the binding buffer was Tris-HCl (pH 8.0) 20 mM, KCl 40 mM, MgCl_2_ 2 mM, ZnCl_2_ 2 μM, 2% (wt/vol) milk, 51.3 ng/μl salmon testes DNA, 0.2 μg/μl bovine serum albumin, 1 mM dithiothreitol.

Pre-incubated protein binding mixtures were applied to individual chambers and incubated for 1 h with the double-stranded DNA chip. The chips were washed once with PBS / 0.5% (v/v) Tween-20 for 3 min and then once with PBS / 0.01% Triton X-100 for 2 min. After the protein incubation and washing steps, Alexa647-conjugated GST antibody (Cell Signaling Technology, Catalog #3445; dilution 1:30), Alexa488-conjugated GST antibody (Invitrogen, Catalog #: A-11131; dilution 1:30); Penta·His Alexa647-conjugated antibody (Qiagen, Catalog #: 35370; dilution 1:20), Penta·His Alexa488-conjugated antibody (Qiagen, Catalog #: 35310; dilution 1:20) in the protein binding buffer / 2% milk were applied on the chip for 1 h at room temperature. Max protein was fluorescently tagged with TAMRA, so no antibody was used. The chips were washed twice with PBS / 0.05% (v/v) Tween-20 for 3 min and then once with PBS for 2 min. Washing steps after each incubation step were performed in Coplin jars at room temperature, on a shaker at 125 r.p.m. The fluorescent signal (at 635 nm, 532nm or 488nm) of bound TFs for each DNA spot was measured at 2.5 μm resolution using a GenePix 4400A® microarray scanner. Signal intensities were extracted using GenePix Pro 7.0 software, and median pixel intensity was reported for each DNA probe. Multiple replicates (10 to 20 replicates) of each sequence were used to quantitatively compare the binding signals between sequences.

**Universal PBM analysis**

To identify DNA motifs recognized by the TF of interest, we analyzed all possible 8-base sequences (8-mers) or 7-base sequences (7-mers) as previously described^10^. In brief, DNA features were grouped into two sets—those containing the 8-mer/7-mer (foreground) and those without (background). We then compare the top half of signal intensities from both sets using a modified Wilcoxon-Mann-Whitney statistic, which helps identify the most enriched 8-mer/7-mer, termed the "seed" of the motif. Next, we assess the contribution of each nucleotide position within this seed by evaluating all possible nucleotide variants at each position, and the motif was further refined by including gaps at positions with high variability. Finally, we convert the derived motif into a position weight matrix (PWM), allowing for a quantitative representation of the TF's binding specificity.

**Bio-layer interferometry (BLI) measurements**

Bio-layer interferometry (BLI) assays were performed using an Octet Red96e System (ForteBio; Menlo Park, CA) in 96-well plates. Streptavidin Octet biosensors (ForteBio; Menlo Park, CA) were dipped into nuclease-free water (Sartorius) for 10 min to hydrate. To obtain the baseline, the sensors were dipped for 60 sec in the kinetic buffer, before dipping into 200 μl of biotinylated DNA probe (50 nM) in the kinetic buffer for the loading step. The DNA probes were constructed by annealing two strands to form an intact binding site or three strands to form a nicked site. Each oligo was purchased from Integrated DNA Technologies (IDT), reconstituted to 100 μM in Nuclease-Free Duplex Buffer (IDT) and mixed to form 1 μM final concentration with the desired stoichiometry (1:1.2 or 1:1.2:1.2, with the biotinylated strand being 1). The resulting mixture was heated to 95 ºC for 5 min and cooled down on ice at 4 ºC. The loading was manually stopped when the response signal reached 0.4 nM for each DNA probe, to ensure an equal amount of DNA probes were immobilised on each sensor. Sensors were then dipped into the kinetic buffer for 60 sec. Next, the protein association step was performed in kinetic buffer at the indicated concentrations for 600 sec (ETS1) or 800 sec (Sox2) to obtain the association curve. Then, the tips were dipped in kinetic buffer again for 400 sec to obtain the dissociation curve. Sensors were then regenerated by dipping the tips in regeneration buffer (2 M NaCl) for 5 sec and then kinetic buffer for 5 sec, repeated three times. All measurements were carried out at 25 ºC. Data were analysed within the ForteBio Data Analysis software. For ETS1, association and dissociation curved were fitted locally with a 1:1 binding model. For Sox2, association and dissociation curved were fitted globally with a 1:1 binding model. *k_on_*, *k_off_*, kinetic *K_D_* and thermodynamic *K_D_* were obtained and reported.

The kinetic buffer for ETS1 contains PBS, 5 mM MgCl_2_ and 0.05% Tween-20. ETS1 protein used was untagged ETS-DBD (residues 280 to 440, 13.6 μM). The kinetic buffer for Sox2 contains 50 mM HEPES (pH 7.5), 150 mM NaCl, 5 mM MgCl_2_ and 0.05% Tween-20. Sox2 protein used was His-Sox2 as in the PBM experiments.

**Crystallisation and structure determination of TBP-nicked DNA complexes**

TBP-AG_noP: To obtain crystals of the nicked TBP-G_noP DNA with TBP, a DNA site with a 4 bp complementary overhang was used (based on the AMVV DNA site). The DNA sites to generate the duplex were (5´-CTATAAAAGCGC-3´ and 5´-TTTTATAG-3´). This duplex DNA was mixed at a 1:1 stoichiometry with TBP (at 10 mg/mL) and hanging drop vapor diffusion screens (Wizard I-IV) were carried out at room temperature (rt). Crystals were obtained by mixing the protein-DNA complex 1:1 with a crystallization solution consisting of 20% (w/v) PEG 3350, 0.1 M Citric acid (pH 5.0) and 0.2 M sodium citrate. The crystals took several days to grow and contain 4 apo TBP molecules and two protein-DNA complexes in the crystallographic asymmetric unit (ASU). The crystals were cryo-preserved by dipping them in a solution consisting of the crystallization reagent supplemented with 20% (v/v) ethylene glycol for 2 s before plunging them into liquid nitrogen. Data were collected at the advanced light source (ALS) beamline 5.0.2 and processed with XDS^11^. The structure was solved by molecular replacement (MR) using the pdb 6UEP as a search model in Phenix^12^. Multiple rounds of refitting in Coot^13^ and refinement in Phenix^12^ was carried out to convergence. See Extended Data Table 1 for data collection and refinement statistics.

TBP-AG_P: To obtain crystals of the nicked TBP-AG_P DNA with TBP, a DNA site with a 4 bp complementary overhang with a 5´phosphate within the nick was used. The DNA sites to generate the duplex were (5´-GCTATAAAAGCGC-3´ and 5´-P-TTTTATAGC-3´). This duplex DNA was mixed at a 1:1 stoichiometry with TBP (at 10 mg/mL) and hanging drop vapor diffusion was used. Wizard I-IV screens were employed at room temperature. Crystals were obtained by mixing the protein-DNA complex 1:1 with a crystallization solution consisting of 10% (w/v) PEG 8000, 0.1 M CHES pH 9.5. The crystals took several days to grow and contain 6 protein-DNA complexes in the ASU. The crystals were cryo-preserved by dipping them in a solution consisting of the crystallization reagent supplemented with 20% (v/v) ethylene glycol for 2 s before plunging them into liquid nitrogen. Data were collected at the advanced light source (ALS) beamline 5.0.2 and processed with XDS^11^. The structure was solved by molecular replacement (MR) using the TBP-TBP-AG_noP protein-DNA complex as a search model^14^. See Extended Data Table 1 for final data collection and refinement statistics.

TBP-TG_noP: To obtain crystals of the nicked TBP-TG_noP DNA with TBP, the DNA sites used to generate the duplex were (5´-GCTATAAATGCGC-3´and 5´-ATTTATAGC-3´). This duplex DNA was mixed at a 1:1 stoichiometry with TBP (at 10 mg/mL) and hanging drop vapor diffusion was used. Wizard I-IV screens were employed at room temperature. Crystals were obtained by mixing the protein-DNA complex 1:1 with a crystallization solution consisting of 4 M sodium Formate, 0.1 M sodium acetate (pH 3.8). The crystals took several weeks to grow to optimal size and contain 1 protein-DNA complex in the ASU. The crystals were cryo-preserved straight from the drop, plunging them directly into liquid nitrogen. Data were collected at the advanced light source (ALS) beamline 5.0.2 and processed with XDS^11^. The structure was solved by molecular replacement (MR) using the TBP-TBP-AG-noP protein-DNA complex as a search model^13,14^. See Extended Data Table 1 for final data collection and refinement statistics.

TBP-TG_P: To obtain the TBP-TBP-TG_P complex a 4 bp complementary overhang with a 5´ phosphate within the nick was used. The DNA sites to generate the duplex were (5´-GCTATAAAAGCGC-3´ and 5´-P-TTTTATAGC-3´). This duplex DNA was mixed at a 1:1 stoichiometry with TBP (at 10 mg/mL) and hanging drop vapor diffusion was used. Wizard I-IV screens were employed at room temperature to find crystallization conditions. Crystals were obtained by mixing the protein-DNA complex 1:1 with a crystallization solution consisting of 20% (w/v) PEG 3350, 0.1 M Citric acid (pH 5.0) and 0.2 M sodium citrate. The crystals took several days to grow and contain 4 apo TBP and two protein-DNA complexes in the ASU. The crystals were cryo-preserved by dipping them in a solution consisting of the crystallization reagent supplemented with 20% (v/v) ethylene glycol for 2 s before plunging them into liquid nitrogen. Data were collected at the advanced light source (ALS) beamline 5.0.2 and processed with XDS^11^. The structure was solved by molecular replacement (MR) using the TBP-TBP-TG_noP protein-DNA complex as a search model. See Extended Data Table 1 for final data collection and refinement statistics.

**Crystallisation and structure determination of EGR1-nicked DNA complexes**

The duplex DNA sequences used for crystallisation were formed by annealing three strands together. The oligos were purchased as a powder from Integrated DNA Technologies (IDT), each solubilised to 1 mM concentration. The three strands were mixed together with the desired stoichiometry (1:1:1.2, with the shortest strand being 1.2) to a concentration of 220 μM in the buffer consisting of 125 mM bis-trispropane HCl (pH 8.0) and 500 mM NaCl. The mixture was heated to 95 ºC for 5 min, followed by overnight slow cooling to 4 ºC to allow for proper annealing. Next, to obtain EGR1 crystals with various nicked DNA sites, the protein was centrifuged for 10 min at 4 ºC to get rid of any possible precipitation, measured concentration with Thermo Scientific Nanodrop One Microvolume UV-Vis Spectrometer, and mixed with the annealed DNA sites at the desired stoichiometry (1:1.1 ratio for protein:DNA) to a final complex concentration of 100 μM. The resulting mixture was then concentrated around 10-fold using a Vivacon® 500 (2,000 MWCO Hydrosart; Sartorius), to a final complex concentration of 1 mM. The resultant protein-DNA complexes were then used in vapour diffusion crystallisation screens.

EGR1-r7noP: Crystals were grown at 19°C using the hanging drop vapor diffusion method. The well solution contained 0.2 M Sodium chloride, 0.1 M MES (pH 6.0), and 20% (w/v) polyethylene glycol monomethy lether (PEG-MME) 2000. Diffraction data were collected to 1.9 Å resolution at 100 K using an in-house Rigaku liquid-metal-jet (LMJ) X-ray Synergy System with a HyPix Arc 150° detector. EGR1-r7noP crystallised in the C222₁ space group, with one subunit in the asymmetric unit. The structure of the Zif268 protein-DNA complex (PDB code 1AAY)^15^ was used as a model for molecular replacement. See Extended Data Table 2 for final data collection and refinement statistics.

EGR1-ren5P: Crystals were grown at 19°C using the sitting drop vapor diffusion method. The well solution contained 0.1 M Magnesium acetate tetrahydrate, 0.1 M Sodium cacodylate (pH 6.5), and 15% (w/v) PEG 6000. Diffraction data were collected to 2.0 Å resolution. EGR1-ren5P crystallised in the C222₁ space group, with one subunit in the asymmetric unit. EGR1-r7noP structure was used to generate the EGR1-ren5P model for molecular replacement. See Extended Data Table 2 for final data collection and refinement statistics.

EGR1-ren7noP: Crystals were grown at 19°C using the sitting drop vapor diffusion method. The well solution contained 0.1 M Calcium acetate, 0.1 M MES (pH 6.0) and 15% (v/v) PEG 400. Diffraction data were collected to 1.95 Å resolution. EGR1-ren7noP crystallised in the C222₁ space group, with one subunit in the asymmetric unit. The structure of the Zif268 protein-DNA complex (PDB code 1AAY)^15^ was used as a model for molecular replacement. See Extended Data Table 2 for final data collection and refinement statistics.

Model Building and Refinement:

Initial models were iteratively rebuilt and refined using Coot^13^ and Phenix^12^. Model geometry was evaluated using MolProbity^16^.

Data Availability Statement

Atomic coordinates and structure factors for TBP-AG_noP, TBP-AG_P, TBP-TG_noP and TBP-TG_P are deposited in the PDB database under accession numbers 9OWI, 9OWZ, 9OW7 and 9OW8, respectively. Atomic coordinates and structure factors for EGR1-r7noP, EGR1-ren5P, and EGR1-ren7noP are deposited in the PDB database under accession numbers 9RIC, 9RI6, and 9RJ6, respectively.

**Structural analysis of Watson-Crisk and nicked DNA structures**

X-ray crystal structures and NMR structures for the desired canonical DNA-protein complexes were downloaded with their PDB information including resolution, macromolecule type etc. from the RCSB webserver. Structures were parsed using X3DNA-DSSR^17^ and Curves+ web server^18^, the structural parameters were extracted and plotted for the purpose of comparison.

The structural analysis of protein-nicked DNA complex was done on the X-ray crystal structures we solved. In X3DNA-DSSR, the data was input as it was in the PDB file. In Curves+ web server, the DNA structure was input as two strands, ignoring the nick. It was only done for the structures with 5´ phosphate at the breakage site, to ensure a well-defined helical backbone.

**Titles and Legends for Supplementary Tables 1-7**

**Supplementary Table 1. PIC-NIC data.**

This file contains the raw PIC-NIC data for the 15 TFs.

**Supplementary Table 2. Structural parameters and contact information.**

This table contains structural parameters generated by x3DNA and H bond contacts defined by ChimeraX at every position in the DNA binding sites for the 15 TFs.

**Supplementary Table 3. Universal PBM data.**

This table contains universal protein binding microarray data for the 15 TFs.

**Supplementary Table 4. Mutation profiles.**

This table contains processed mutation profile data for the 15 TFs.

**Supplementary Table 5. Universal PBM data for ETS1 mutants.**

This table contains universal protein binding microarray data for the four ETS1 mutants with replicates.

**Supplementary Table 6. BLI data.**

This table contains the raw and fitted data of BLI experiments for ETS1 and SOX2.

**Supplementary Table 7. Structural parameters of TBP/nicked DNA complexes.**

This table contains structural parameters generated by x3DNA for obtained high-resolution crystal structures of TBP/nicked DNA complexes.

**Supplementary Table 8. The composition of the molecular simulation box.**

This table contains the number of molecules in each of the simulation box based on the topology file.
